# Data-Driven Classification of Spectral Profiles Reveals Brain Region-Specific Plasticity

**DOI:** 10.1101/782979

**Authors:** Christina Lubinus, Joan Orpella, Anne Keitel, Helene Gudi-Mindermann, Andreas K. Engel, Brigitte Roeder, Johanna M. Rimmele

**Affiliations:** Department of Neuroscience, Max-Planck-Institute for Empirical Aesthetics, Grüneburgweg 14, D - 60322 Frankfurt am Main, Germany; Department of Psychology, New York University, 6 Washington Place, New York, NY 10003, USA; Psychology University of Dundee, Scrymgeour Building, Dundee DD1 4HN, Scotland, UK; Biological Psychology and Neuropsychology, University of Hamburg, Von-Melle-Park 11, D - 20146 Hamburg, Germany; Deparartment of Social Epidemiology, University of Bremen, Grazer Straße 2, D - 28359 Bremen, Germany; Department of Neurophysiology and Pathophysiology, University Medical Center Hamburg-Eppendorf, Martinistraße 52, D - 20246 Hamburg, Germany

**Author notes:** <u>Corresponding author:</u> Johanna M. Rimmele, PhD, Department of Neuroscience, Max-Planck-Institute for Empirical Aesthetics, Grüneburgweg 14, D - 60322 Frankfurt am Main, Germany.

**Keywords:** congenital blindness, MEG, oscillations, spectral fingerprints

## Abstract

Congenital blindness has been shown to result in behavioral adaptation and neuronal reorganization, but the underlying neuronal mechanisms are largely unknown. Brain rhythms are characteristic for anatomically defined brain regions and provide a putative mechanistic link to cognitive processes. In a novel approach, using magnetoencephalography resting state data of congenitally blind and sighted humans, deprivation-related changes in spectral profiles were mapped to the cortex using clustering and classification procedures. Altered spectral profiles in visual areas suggest changes in visual alpha-gamma band inhibitory-excitatory circuits. Remarkably, spectral profiles were also altered in auditory and right frontal areas showing increased power in theta-to-beta frequency bands in blind compared to sighted individuals, possibly related to adaptive auditory and higher-cognitive processing. Moreover, occipital alpha correlated with microstructural white matter properties extending bilaterally across posterior parts of the brain. We provide evidence that visual deprivation selectively modulates spectral profiles, possibly reflecting structural and functional adaptation.

## Introduction

Brain rhythms occur ubiquitously across cortex (Buzsáki 2004; Buzsáki et al. 2013; Singer 2018) and have been related to cognitive functions. Investigating changes in brain rhythms in congenital blindness can provide crucial insights into the neuronal mechanisms of behavioral changes as found in this population. Congenitally blind individuals perform better on various auditory, tactile, and higher-cognitive tasks compared to sighted controls (Bull et al. 1983; Lessard et al. 1998; Roeder et al. 2001; Amedi et al. 2003; Gougoux et al. 2005; Foecker et al. 2012), including tasks where certain aspects of temporal processing are essential, such as in temporal order processing tasks, task involving musical meter or ultra-fast speech processing (Hoetting et al. 2004; Roeder et al. 2004; Stevens and Weaver 2005; Moos and Trouvain 2007; Roeder et al. 2007; Trouvain 2007; Hertrich et al. 2009; Dietrich et al. 2013; Lerens et al. 2014; Carrara-Augustenborg and Schultz 2019; Zhang et al. 2019). Improved behavioral performance in auditory or tactile tasks in congenitally blind individuals is accompanied by cross-modal plasticity. That is, the “visual” cortex has been observed to be recruited during non-visual tasks (Burton 2003; Pascual-Leone et al. 2005; Bedny et al. 2011; Voss and Zatorre 2012; Gudi-Mindermann et al. 2018; Rimmele et al. 2019), which has been postulated to functionally contribute to compensatory task performance. Additionally, intramodal plasticity has been observed in congenitally blind individuals. That is, the auditory (Elbert et al. 2002; Stevens and Weaver 2005; Gougoux et al. 2009; Schepers et al. 2012; Hertrich et al. 2013; Watkins et al. 2013) and somatosensory (Pascual-Leone and Torres 1993; Roeder et al. 1996; Sterr et al. 1998a; Sterr et al. 1998b) cortices were reported to show altered (mostly increased) neuronal activity during task processing. Intramodal plasticity has been related to early postnatal sensory experiences shaping the functional development of cortical representations as shown in animal research (Zhang et al. 2001; for review: Rauschecker 2008). For instance, reorganization of primary somatosensory and auditory areas has been reported in cats that were visually deprived from birth (Rauschecker et al. 1992; Korte and Rauschecker 1993). Although these behavioral and neuronal adaptations have been observed, the mechanisms underlying visual deprivation-related plasticity are still poorly understood.

Studying brain rhythms in congenitally blind individuals might provide further insights into the mechanisms of neuronal plasticity, as brain rhythms reflect neuronal circuitry activity and have been related to various sensory, motor and higher-cognitive processes, as discussed in the following. Brain rhythms are thought to reflect the synchronization (phase-alignment) of oscillatory activity across neuronal populations, subserving the formation of both local assemblies and large-scale functional networks (Engel et al. 1992; Hipp et al. 2011; Raichle 2011; Siegel et al. 2012; Singer 2013). Specific spectral profiles have been conjectured to be associated with different anatomical areas (Giraud et al. 2007; Keitel and Gross 2016). Such spectral profiles are assumed to reflect the intrinsic brain rhythms of a brain area that are related to distinct processes (Keitel and Gross 2016). Importantly, intrinsic brain rhythms measured spontaneously (e.g., ongoing activity during resting state) reflect the functional organization of the brain and thus the default architecture recruited during task-related processes (Smith et al. 2009; Deco et al. 2011; Raichle 2011; Engel et al. 2013; Raichle 2015; Keitel and Gross 2016; Sormaz et al. 2018) because intrinsic brain rhythms oscillate in a characteristic way even when not involved in task-related processing (Brookes et al. 2011; Deco et al. 2011). For example, Keitel and Gross (2016) have shown that a characteristic pattern of delta-, theta-, alpha-, and beta-band activity is present in primary auditory cortex both during rest and speech comprehension. While the global appearance of the spectral profile of primary auditory cortex was similar during speech comprehension and rest, the relative contribution of individual frequency clusters (i.e., their spectral power) was altered by the condition. During speech comprehension, in comparison to rest, the alpha band was suppressed, indicating reduced inhibition of sensory cortices (Jensen and Mazaheri 2010). The other bands, which were associated with several levels of speech processing, were enhanced.

Various studies have observed that brain rhythms are recruited in a task-specific manner during sensory, motor and higher-cognitive tasks, linking intrinsic brain rhythms to cognitive functions (Engel et al. 1992; Singer and Gray 1995; Engel et al. 2001; Lakatos et al. 2008; Holcombe 2009; Giraud and Poeppel 2012; Lakatos et al. 2013; Heusser et al. 2016; Portoles et al. 2018; Rimmele, Gross, et al. 2018; Schroeder et al. 2018; VanRullen 2018). For instance, in sighted individuals, the role of the alpha rhythm (~8–12Hz) in cognition is well-established, with posterior alpha- and gamma-band oscillations coupling in excitatory-inhibitory cycles (Klimesch et al. 2007; Buffalo et al. 2011; Jensen et al. 2012). Specifically, the alpha rhythm mediates the inhibition of task-irrelevant neuronal circuits. Gamma-band oscillations (30–100 Hz) in occipital cortex, by contrast, allow for feedforward processing of task-relevant visual information (van Kerkoerle et al. 2014; Michalareas et al. 2016). In congenitally blind individuals, a reduced or absent occipital alpha rhythm has repeatedly been observed (Adrian and Matthews 1934; Noebels et al. 1978; Kriegseis et al. 2006; Hawellek et al. 2013; Schubert et al. 2015). The lack of visual input during a sensitive period of brain development in congenitally blind individuals presumably results in atrophy or reorganization of posterior alpha generators (Ptito et al. 2008; Wang et al. 2013; Aguirre et al. 2016; Reislev et al. 2016). Posterior alpha generators have been located within the cortical and thalamo-cortical visual pathways (Lopes da Silva 1991; Lőrincz et al. 2009) and posterior gamma generators within the cortical visual pathways (Bastos et al. 2014; Marshall et al. 2018). Beyond the occipital brain rhythms, however, no clear hypothesis linking brain rhythms and structural alterations in congenitally blind individuals can be made.

In sighted individuals, beyond the alpha rhythm, intrinsic theta-band oscillations (~4–7 Hz) in auditory cortex (Lakatos et al. 2005; Keitel and Gross 2016) have been widely studied. Theta brain rhythms likely play a central role in the temporal segmentation of speech signals during speech comprehension (Luo and Poeppel 2007; Giraud and Poeppel 2012; Gross et al. 2013; Doelling et al. 2014; Pittman-Polletta et al. 2020), as well as segmentation of music and environmental sounds (Henry et al. 2014; Doelling and Poeppel 2015). Accordingly, initial findings have linked the ability of congenitally blind individuals to comprehend ultra-fast speech–at rates where speech comprehension fails in sighted individuals–to accelerated theta brain rhythms in auditory cortex, as well as the recruitment of additional brain areas such as the visual cortex (Trouvain 2007; Hertrich et al. 2009; Dietrich et al. 2013; Hertrich et al. 2013; see also: Van Ackeren et al. 2018). Notably, functional connectivity measures suggest that brain rhythms in the beta-and gamma-band in brain areas outside of visual cortex might also be altered in congenitally blind compared to sighted individuals (Schepers et al. 2012; Gudi-Mindermann et al. 2018; Rimmele et al. 2019). However, as these alterations are not well understood, we refrain from discussing them here. In summary, the systematics of how spectral profiles outside of visual cortex are altered due to congenital visual deprivation, are not well understood and no clear hypothesis can be derived beyond the occipital alpha rhythm.

The goal of the present study is to characterize alterations in spectral profiles in congenitally blind as compared to sighted individuals in order to map visual deprivation-related spectral changes to brain areas. By studying a broad range of frequencies, instead of narrow frequency bands, our research provides the means towards linking changes in spectral profiles across the whole brain to adaptive behavior in congenitally blind individuals in future research. Standard analyses of brain rhythms face several problems (see Keitel and Gross 2016), which possibly explains the lack of a comprehensive account of spectral changes in congenitally blind individuals beyond the alpha rhythm. Predominant neuronal activity in the alpha and super-low frequency ranges (1/f) complicates the analysis of spectral properties in the lower frequency ranges. Furthermore, the brains’ temporal dynamics are often captured poorly over the course of the recording session (de Pasquale et al. 2010; Singer 2013). Here, we have employed a novel analysis pipeline, introduced by Keitel and Gross (2016), which reveals brain area-specific spectral profiles. The analysis pipeline overcomes the limitations of standard analyses of brain rhythms by using segment-based clustering (of source-localized Fourier spectra) to disentangle spectral properties in the lower frequency ranges and by taking the time course of activity into account. We extended the pipeline (Fig. 1) to quantify differences in spectral profiles across cortical brain areas between congenitally blind and sighted individuals by using a cross-classification approach. This approach allowed us to identify brain areas with changed spectral profiles as a result of congenital blindness. More specifically, the clustering procedure reveals multi-dimensional spectral profiles that might consist of several clusters reflecting distinct intrinsic brain rhythms.

**Figure 1.**
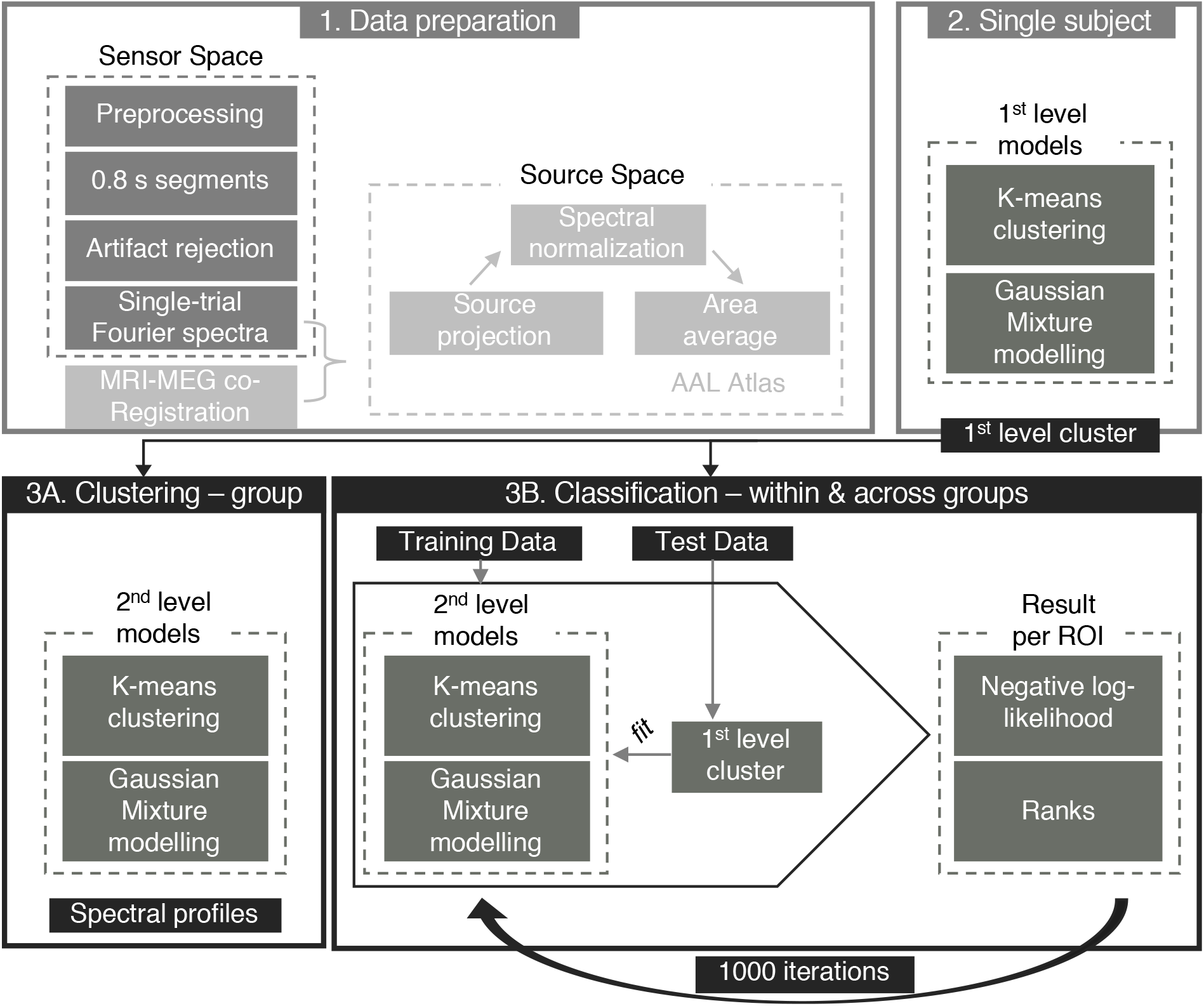
Analysis pipeline adapted from Keitel and Gross. (2016).(1) Continuous resting state MEG data were preprocessed and segmented into trials of 0.8 s length. Complex Fourier spectra were computed for each trial separately and projected into source space using previously defined beamforming (LCVM) coefficients. The data were spatially normalized, dividing each voxel’s power by the mean power of all trials and voxels. Voxels were grouped according to the AAL atlas and power values were averaged across voxels of each anatomical area (N = 115). (2) In the 1^st^-level analysis, power matrices were clustered into 9 distinct spectral clusters per participant and brain region using k-means and Gaussian Mixture model algorithms. (3A) In the 2^nd^-level analysis, 1^st^-level individual clusters were again subjected to k-means clustering and GMMs to establish region-specific spectral clusters consistent at the group level, also referred to as spectral fingerprints. The optimal number of group-level clusters per anatomical area was defined by the Silhouette Criterion prior to the group-level clustering procedure. (3B) For the classification procedure the experimental group was divided into training and test set. For each brain region, the fit between 1^st^-level clusters of the test group and group-level clusters of all regions of the training set was calculated. For each anatomical region, this resulted in a negative log-likelihood value for all regions (i.e., 115 values per region), indicating its similarity to all brain regions based on the spectral clusters. This fitting procedure between training and test set was repeated 1000 times with new group assignment (training vs. test) on each iteration. The interindividual variance within test and training groups of each iteration was controlled for with an additional 100 iterations within the respective sets.

First, we expected to replicate Keitel and Gross (2016) by showing that spectral profiles in sighted adults are region-specific, enabling the classification of brain regions based on spectral profiles. Second, we hypothesized that within a group of congenitally blind individuals, spectral profiles would similarly follow specific patterns and enable the classification of brain regions. Third, visual deprivation-related plasticity was predicted to result in altered spectral profiles in congenitally blind individuals as compared to the sighted, particularly for brain regions for which visual deprivation-related reorganization (cross- and intra-modal plasticity) has previously been shown, such as for sensory cortices. A specific hypothesis regarding the changes in brain rhythms in congenitally blind individuals was only derived for the alpha rhythm, which was expected to be reduced in occipital cortex. Finally, we expected a relationship of microstructural white matter properties and spectral profiles of occipital cortex.

## Materials and Methods

### Participants

The study was approved by the German Psychological Association. All participants gave written informed consent prior to the experiments and received monetary compensation. The data were recorded in the context of a larger project (Gudi-Mindermann et al. 2018; Rimmele et al. 2019). Three to four minutes of resting state MEG data were collected from a group of sighted and congenitally blind individuals matched in age, gender and education. During data collection the congenitally blind (CB) and the sighted (S-BF) participants were blindfolded. The blindfolding of CB participants aimed at reducing involuntary eye movements. Additionally, for the sighted a resting state measurement with open eyes was conducted (S-EO). The data reported here include (after a few subjects were excluded, see below) 26 subjects for the CB (12 females; mean age: 37.8 years; *SD*: 10.2 years; age range: 22–55 years), 24 for the S-BF (11 females; mean age: 36.8 years; *SD*: 10.1 years; age range: 21–55 years) and 23 for the S-EO (11 females; mean age: 37.3 years; *SD*: 9.8 years; age range: 21–55 years). A few subjects were excluded after data collection because of corrupted resting state files (one subject for the CB, one subject for the S-EO) or lack of individual structural MRI scan (three subjects for the S-BF and S-EO). All participants were healthy with normal hearing (self-report) and assured no history of psychiatric or neurological disorders. One blind participant reported a history of a depressive mood disorder, but was free of symptoms and without current treatment. Sighted participants had normal or corrected-to-normal vision (self-report). In the blind, vision loss was total and resulted from a variety of peripheral (pre)natal conditions (retinopathy of prematurity: *n*=9; genetic defect, *n*=5; congenital optic atrophy: *n*=2; Leber's congenital amaurosis: *n*=2; congenital cataracts, glaucoma: *n*=2; congenital retinitis: *n*=2; binocular anophthalmia: *n*=2; retinitis pigmentosa: *n*=1; congenital degeneration of the retina, *n*=1). Seventeen participants reported minimal residual light perception.

### MRI and MEG Data Acquisition

For all participants T1-weighted structural MRI scans and DWI-MRI scans were obtained with a 3T scanner (Siemens Magnetom Trio, Siemens, Erlangen, Germany). For the T1-weighted images we used the following parameters: TE = 2.98 ms, TR = 2300 ms, flip angle = 9, and isotropic 1 mm3 voxels, 256 sagittal slices. The MEG data were recorded in a magnetically shielded room using a 275-channel whole-head system (Omega, 2000, CTF Systems Inc.), while participants sat in an upright position. The data were acquired with a sampling rate of 1200 Hz. Prior to each experiment, the head position was measured relative to the MEG sensors and during the recording the head position was tracked.

### Data Analysis

The initial analyses in this study are adopted from the analysis pipeline proposed by Keitel and Gross (2016). The modifications to the analysis pipeline and the novel analyses are stated in detail below. All analyses were carried out using Matlab R2018a (version 9.4.0.813654, The Math Works, Inc.), the Fieldtrip Toolbox (version 20181104), and SPM12.

### Data Preparation in Sensor Space: Preprocessing, Artifact Rejection, Source Localization

During preprocessing, the MEG signal was downsampled to 250 Hz, denoised and detrended. To better capture the dynamically changing spectral properties of the brain, the continuous signal was segmented into trials of 0.8 s. Trials were declared as noisy and excluded when their zscore was higher than 2. On average, seven trials were excluded, resulting in a mean of 340.3 trials (*SD*= 34.7) per subject (S-EO: mean = 346.7, *SD* = 37.4; S-BF: mean = 336.8, *SD* = 34.2; CB: mean = 338, *SD* 33.2). Due to shorter recordings in the present study, trial duration was slightly shorter, relative to the 1 s duration used in Keitel and Gross (2016), to increase the amount of trials. MEG channels were labeled as noisy and rejected when the ratio between their noise level (in *SD*) and that of the neighboring sensors (in *SD*) exceeded a value of 0.5 ([Sensor *SD* – Neighbor *SD*] / Neighbor *SD*; mean number of excluded channels = 1.22, *SD* = 1.34). Finally, using independent component analysis (ICA), data was cleaned from heartbeat, eye blinks and eye movements-related artifacts (components were identified based on their timecourse, topography and variance across trials). To prepare the source projection of the Fourier spectra, beamformer coefficients were obtained. For this purpose, we applied co-registration of individual T1-weighted MRI scans and the MEG coordinate system, realignment, segmentation and normalization to Montreal Neurological Institute (MNI) space. A forward model was created using a single-shell model and linearly constrained minimum variance (LCMV) beamformer coefficients (Van Veen et al. 1997) were calculated for the MEG time series for each individual voxel on the 10 mm regular grid.

### Spectral Analysis in Sensor Space

The analyses described in the following were performed for all three groups separately (CB, S-EO, S-BF; for an overview, see Fig. 1). First, Fourier spectra were calculated on 0.8 s long trials for each subject, using a multitaper approach (3 tapers) and zero-padding (length of 2 s). We slightly reduced the duration of the temporal segments compared to the duration used by Keitel and Gross (2016), i.e., 1 s, in order to match the overall number of segments used for the analyses while accounting for the shorter overall MEG recording time in our study. Second, using the previously computed LCMV coefficients, the complex Fourier spectra were projected into source space. Neuronal power, i.e., the squared amplitude of the Fourier coefficient, was computed for each time segment, voxel, and frequency. Furthermore, neuronal power of individual voxels and segments was ratio normalized, i.e., divided by the mean power across all voxels and trials. This ratio normalization resulted in voxel-specific spectral properties with values above/below 1 highlighting the differences of a given voxel to the mean spectral power across all voxels separately at each frequency. All values were subtracted by 1 (leading to values above/below zero), to facilitate the identification of changes in power (decreases/increases). We additionally performed a control analysis without the normalization (Fig. S5). This analysis confirms our findings, while it demonstrates the advantage of the normalization procedure.

### k-Means Clustering and Gaussian Mixture Modelling of Source-Localized Spectral Activity

To identify region-specific spectral clusters in the individual subject, the brain was parcellated according to the AAL atlas (Tzourio-Mazoyer et al. 2002; 116 regions of interest, ROIs). For one anatomical region (cerebellum 3L), however, the interpolation between the AAL atlas and the source model was not successful. Thus, this region was excluded and all analyses are based on the remaining 115 anatomical areas. For each of the ROIs, voxels were grouped and power spectra were averaged across voxels. Within a brain area, neuronal populations might express several distinct intrinsic brain rhythms, which together constitute the spectral profile of an area. Clustering algorithms were employed to identify spectral clusters, reflecting intrinsic brain rhythms. First, trial-by-frequency matrices were subjected to a k-means algorithm (MacQueen 1967) which established spectral clusters by partitioning the *n* observations (0.8 s temporal segments) into *k* clusters. For the 1^st^-level analysis, the *k* was set to 9, based on the Silhouette criterion evaluation (Rousseeuw 1987). Second, for each subject and ROI, Gaussian Mixture Models (GMMs; Reynolds and Rose 1995) were fitted to the 9 clusters obtained from the kmeans analysis (1^st^-level GMM). Next, in order to identify the optimal number of clusters per brain region across all subjects for the 2^nd^-level group analysis, the 1^st^-level GMMs were evaluated using the Silhouette criterion. Silhouette values were computed for cluster solutions in the range from 1 to 15, the fitting was repeated 1000 times. At the group level, k-means clustering was applied to the 1^st^-level clusters in order to disclose consistent patterns across subjects. The optimal number of clusters per brain area, as assessed by the Silhouette criterion evaluation (Rousseeuw 1987), was used as k-parameter for the algorithm. As before, k-means results were fed into GMM revealing the final clusters per brain region (2^nd^-level GMM).

Clusters were considered for visualization only if they were reflective of the majority of participants. To facilitate reading of the spectral plots, group-level clusters were color-coded according to the frequency of the maximum amplitude of the cluster (peak frequency) (delta: 1–3.5 Hz, red; theta: 4–8 Hz, green; alpha: 8.5–12.5 Hz, blue; beta: 14–30.5 Hz, yellow; gamma: 33.5–100 Hz, magenta). Furthermore, we computed the relevance of each cluster per brain region by analyzing the amount of single subject trials during which a cluster was present. Group clusters (Fig. 1, step 3) were traced back to single subject clusters and the amount of trials that contributed to a single subject cluster (Fig. 1, step 2) was calculated and expressed as percentage. Percentages were averaged across subjects.

### Automatic Within-Group Classification

A classifier was employed to test the specificity of region-specific spectral fingerprints. After splitting each group into half (training and test group), group-level clusters were calculated for the training group for all anatomical regions using k-means and GMM clustering. For each brain region and participant of the test group, the similarity of spectral profiles was assessed as compared to all brain regions of the 2^nd^-level group clusters of the training group by computing the negative log-likelihood for all pairs of regions. This procedure, i.e., group assignment and classification, was repeated 1000 times (note that for the S-EO one subject was left out in every iteration to yield an even number of participants in training and test groups). On each iteration, an additional loop (*N* = 100) controlled for interindividual noise within a group by randomly drawing the adequate number of subjects (i.e., *N*_*S-EO*_ = 11, *N*_*S-BF*_ = 12, *N*_*CB*_ = 13) from the group with replacement, allowing a subject to enter multiple times or not at all. Put differently, within one iteration (*N* = 1000) each participant belonged to either the training or the test group. To account for individual differences, the group clusters were calculated 100 times with a different subset chosen from the respective group each time and finally averaged to obtain a robust group estimate. Based on the mode of clusters identified per brain region in the 2^nd^-level cluster analysis, the optimal number of clusters for the classification analysis was *k* = 2. Likelihood values were ranked and averaged across iterations (20% trimmed mean). For further comparisons, only corresponding ROIs (e.g., how is the Heschl ROI in the test set ranked based on the training set Heschl ROI) were considered.

In addition to the descriptive report of the classification performance, here we tested whether a specific ROI (of the test set) was classified significantly better by the corresponding area of the training set as compared to all other 115 ROIs. This allowed us to exclude the possibility that classification performance was caused by unspecific effects—that is, generic fingerprints. To this end, each region’s mean rank (averaged across iterations) was tested against a distribution of classification ranks generated from all other ROIs (null distribution).

### Automatic Cross-Group Classification

In order to identify differences in region-specific spectral properties between the CB and S-BF, we performed a cross-group classification. We employed the same classification procedure as before; however, the classifier was trained on one group (S-BF), while the other (CB) was utilized as the test set. As before, the classification procedure was repeated 1000 times, drawing a subset of *N* = 12 per group on every iteration. The randomization of subjects chosen on each iteration was identical to the one used for the within-group classification in the S-BF (this is the reason why *N* = 12, instead of using all subjects of both groups). Thus, differences in the classification, as reflected by the ranks, could not be caused by the training set per se. In order to understand whether some brain areas in the CB were not classified well based on the S-BF spectral profiles (i.e. whether the classification of brain regions was different in the cross-group condition as compared to the within-S-BF classification), we tested the cross-group classification mean ranks against the distribution of ranks from the same ROI from the S-BF group (here the null distribution). The distributions were generated by taking the classification rank of a corresponding area from training and test set (i.e. Calcarine) across all iterations (see Fig. S2 for the distributions of all brain areas). We calculated the 95^th^ percentile of the distribution and tested whether the cross-group mean rank of the current region fell above (significant) or below (not significant) this threshold.

To further assess the spectral profiles of brain areas that were significantly different in the cross-group classification, post-hoc permutation statistics were applied to the raw, normalized region-specific spectra (i.e., Fourier spectra without clustering procedure). This additional analysis was performed to evaluate the cross-group differences between the sighted and the congenitally blind groups by using a simpler, more established approach. The analysis confirmed the findings from the spectral clustering approach. The spectral analysis was calculated as in the main analysis (see above). For all significant brain regions separately, power was averaged across voxels and segments, resulting in a single power value per frequency and per subject. Based on frequency by subject matrices for the CB and the S-BF, group differences in spectral power were tested against a distribution where the group assignment (CB vs. S-BF) was randomly permuted (*N* = 1000). To control for multiple comparisons, we used FDR (Q = 0.05).

### Microstructural White Matter Properties

DW-MRI data were acquired together with T1-weighted structural scans described above. We used an echo planar imaging (EPI) sequence optimized for DWI-MRI of white matter covering the whole brain (64 axial slices; bandwidth = 1502 Hz/Px, 104 × 128 matrix, TR, 8,200 ms; TE, 93 ms; flip angle, 90°; slice thickness, 2 mm; voxel size, 2 × 2 × 2 mm3). The protocol comprised three acquisitions yielding a total acquisition time of 9 min 51 s. This resulted in a total of 120 diffusion-weighted volumes with six interleaved non-diffusion-weighted volumes (b values of 1,500 s/mm2). DWI-MRI scans were acquired from a subset of the original sample of 16 blind and 12 sighted participants.

### DTI-MRI preprocessing and analysis

Diffusion data processing initially corrected for eddy current distortions and head motion by using FMRIB’s Diffusion Toolbox (FDT; FMRIB Software Library; FSL 5.0.1; http://www.fmrib.ox.ac.uk/fsl/; Jenkinson et al. 2012). For a more accurate estimate of diffusion tensor orientations, the gradient matrix was rotated to correct for head movement, using the fdt_rotate_bvecs program in FSL. We then used the Brain Extraction Tool (Smith 2002) for brain extraction, also part of the FSL distribution. Analysis continued with the reconstruction of the diffusion tensors using FSL’s DTIFIT program. FA and RD maps for each participant were calculated using the eigenvalues extracted from the diffusion tensors. Note that FA maps are required in the early stages of TBSS to compute the registrations to MNI standard space and subsequently create the diffusion skeletons. However, we focused our analysis on RD, as this is a more specific measure of diffusivity in white matter than FA or mean diffusivity. Indeed, although several factors can contribute to producing particular RD values, including the number of axons and axon packing and diameter, RD has been most consistently related to myelin content along axons, with increased RD values reflecting higher demyelination (Song et al. 2002; Song et al. 2005; Klawiter et al. 2011; Zatorre et al. 2012). In animal studies, directional measures such as RD, unlike summary parameters such as mean diffusivity or FA, provide better structural details of the state of the axons and myelin (Aung et al. 2013).

Voxel-based analyses of RD maps were performed with TBSS (Smith et al. 2006). Participants’ FA maps (necessary to calculate the registrations to MNI standard space and create the RD skeletons) were registered to the FMRIB58_FA template (MNI152 space and 1 × 1 × 1 mm3) using the nonlinear registration tool (Andersson et al. 2007). These registered FA maps were first averaged to create a mean FA volume. A mean FA skeleton was then produced, representing the centers of all white matter tracts common to all participants in the study. Each participant’s aligned FA data were then projected onto this skeleton by searching for the highest FA value within a search space perpendicular to each voxel of the mean skeleton. This process was repeated for the RD maps by applying the transformations previously calculated with the FA maps. This resulted in individual RD skeletons for each participant. Finally, to assess white matter differences between CB and sighted participants, independent-samples *t*-tests were performed on the RD skeleton. Significant results are reported at FWE-corrected *p* < 0.05 using threshold-free cluster enhancement (Smith and Nichols 2009) and a nonparametric permutation test with 5,000 permutations (Nichols and Holmes 2002). Significant cluster results were averaged and a mean value per participant, reflecting individual microstructural differences, was obtained.

For the mapping between RD values and the standard probabilistic atlases of white matter pathways, all voxels that differed significantly in RD values between the CB and sighted were included. We report only tracts that showed an overlap with these voxels, with the tracts from the probabilistic atlas thresholded at 0.95 probability.

Spearman correlations were used to analyze the correlation between RD values (across groups) and the spectral profiles of brain areas in the occipital cortex. For these areas, peak power values were retrieved for all clusters (i.e., lines showing clear power peak) of the complex spectral profile for each subject, and correlated with the RD values. Multiple comparisons were controlled for by using FDR (Q = 0.05).

## Results

### Spectral Fingerprints Replicate

In our sample of sighted adults with open eyes we successfully replicated the classification of individual brain regions by their spectral profiles as first reported by Keitel and Gross (2016). Particularly, for all tested 115 ROIs the mean classification performance, indicated by the classification ranks, was high (Fig. 2A). Classification ranks refer to the probability of a region being identified by the classifier: For example a mean rank of 1 indicates that a region was correctly assigned (i.e., highest probability among all areas) on every iteration; a mean rank of 2 means that the assignment was correct in many but not all of the iterations (i.e., had the second highest probability among all areas). Here, the mean rank (averaged across all iterations and brain areas) obtained from the classifier analysis was 2.70 (range of ranks: 1–12.7, Keitel mean rank = 1.8), or 2.32 when considering identification of the homologue (left/right hemisphere) areas as a hit (Keitel homologue mean rank = 1.4). Mean ranks of all ROIs are depicted in the histogram and surface plot in Fig. 2A. We statistically quantified the classification performance using permutation tests. The mean classification rank of an area (e.g., right calcarine) was tested against a distribution of classification ranks of all brain areas (except the current one, e.g., right calcarine) accumulated across all iterations (*N* = 1000). For an area with a characteristic spectral profile, classification between corresponding areas (e.g., right calcarine in training vs test set) should be best and thus fall above the 95^th^ percentile of the generated null-distribution. This analysis revealed that for 97% of all areas classification was significantly better when identifying themselves compared to all other regions.

**Figure 2.**
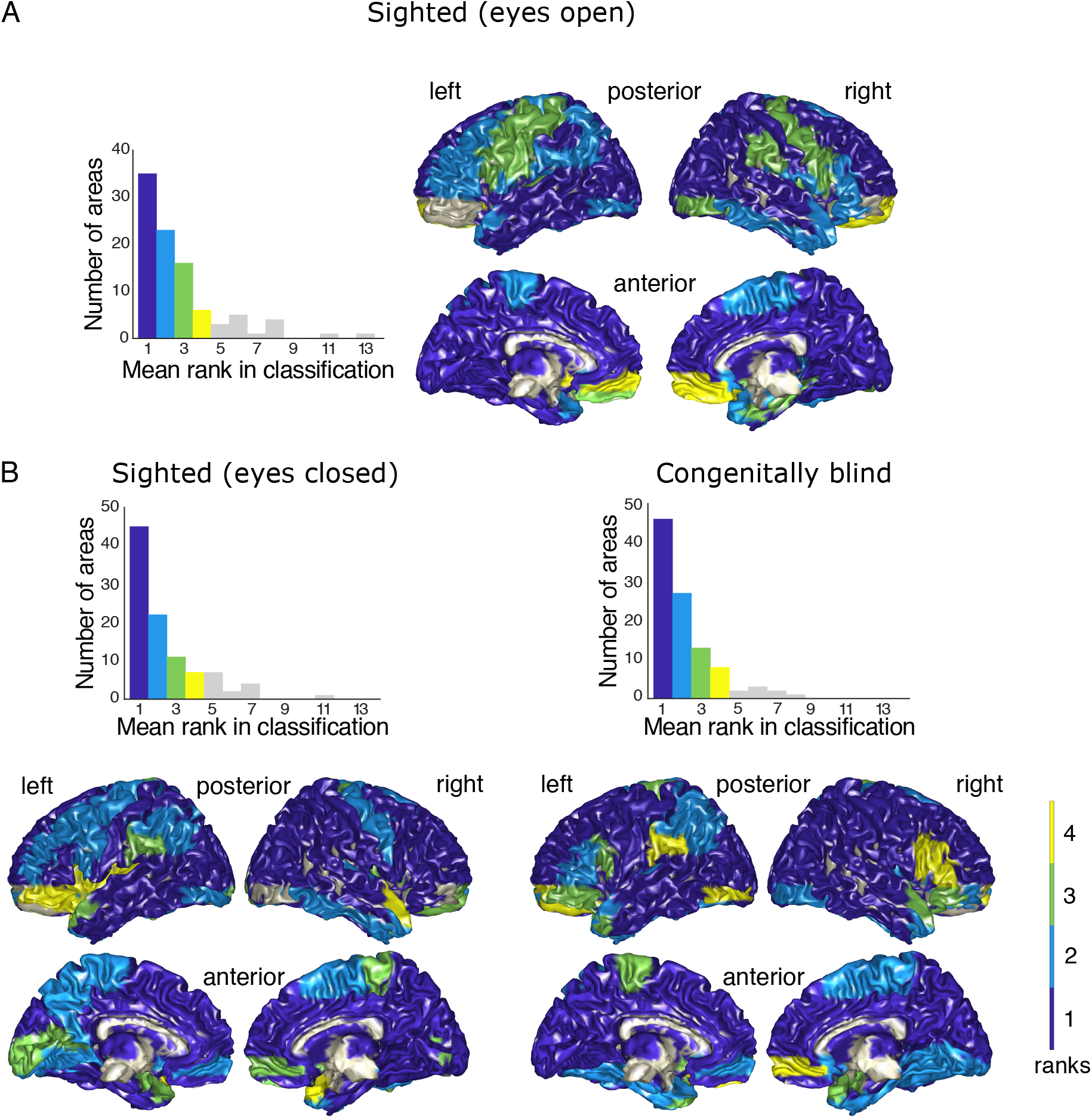
Classification results for all experimental groups. **(A) Sighted group with open eyes (replication sample)**. *Left*. Histogram of mean ranks in classification across all 115 brain areas. 84.5% of the brain regions obtain a mean rank between 1 and 4, while 15.5% of regions were assigned ranks up to 13. (in Keitel and Gross (2016) all mean ranks were between 1 and 4). *Right.* Topography of mean ranks (colors match ranks from the histogram). **(B) Sighted group with eyes closed (*left*) and group of congenitally blind individuals (*right*)**. For both groups, the histogram of mean ranks in classification across all 115 brain areas is displayed in the upper panel and the topography of mean ranks (colors match ranks from histogram) is displayed in the lower panel. Bin width is 1 for all subplots.

Furthermore—although the average optimal number of clusters per anatomical area was lower in our sample (3.4 +/− 2.3 clusters per area (see Fig. 5 and Fig. S1, Supplementary Data) vs. 4.1 +/− 1.86 (*M* + *SD*) in Keitel and Gross (2016)—the clustering approach revealed comparable spectral fingerprints between the studies. Interestingly, for deeper subcortical brain structures (e.g., thalamic and limbic areas), the clusters were less characteristic in the present data (i.e., only a few clusters per area with less specific shapes and high classification ranks; see Fig. S1), possibly reflecting limitations of the signal-to-noise ratio of the used MEG system.

### Good Classification Within Sighted and Congenitally Blind Groups

Our second hypothesis stated that, within a group of congenitally blind individuals, anatomical areas are characterized by specific (although possibly altered as compared to the sighted) spectral fingerprints. We performed the classification procedure for the CB and observed good classification ranks (similar to the ones observed for the S-EO; Fig. 2B; mean rank = 2.51, range = 1–10.3, homologue mean rank = 2.10, percent significant ROIs = 100%), indicating consistent spectral clusters of brain areas in congenitally blind participants. The same procedure was performed on the data of the S-BF and revealed similarly good classification ranks (S-BF: mean rank = 2.64, range = 1–11.4, homologue mean rank = 2.17, percent significant ROIs = 98%) as compared to the S-EO and CB.

Ensuring good within-group classification in the CB and the S-BF was an important prerequisite for consecutive between-group analyses as it promised to prevent potential group differences from arising from large within-group variance. Furthermore, the results showed a similar distribution of mean ranks across the cortical surface for both the CB and the S-BF group, such that the majority of brain regions (~70 out of 115) was identified as best or second-best.

### Spectral Changes in Sensory and Right Frontal Regions in Congenitally Blind Individuals

Based on the literature on intra-and cross-modal plasticity in the CB, we hypothesized that spectral properties would differ between the congenitally blind and normally sighted individuals. To test if (and which) brain areas differed in their spectral properties between the two groups, we implemented a cross-group classification drawing samples from the S-BF for the training and samples from the CB for the test group. This analysis resulted in a mean rank of 5.3 (range = 1.09-27.17) (Fig. 4A).

**Figure 4.**
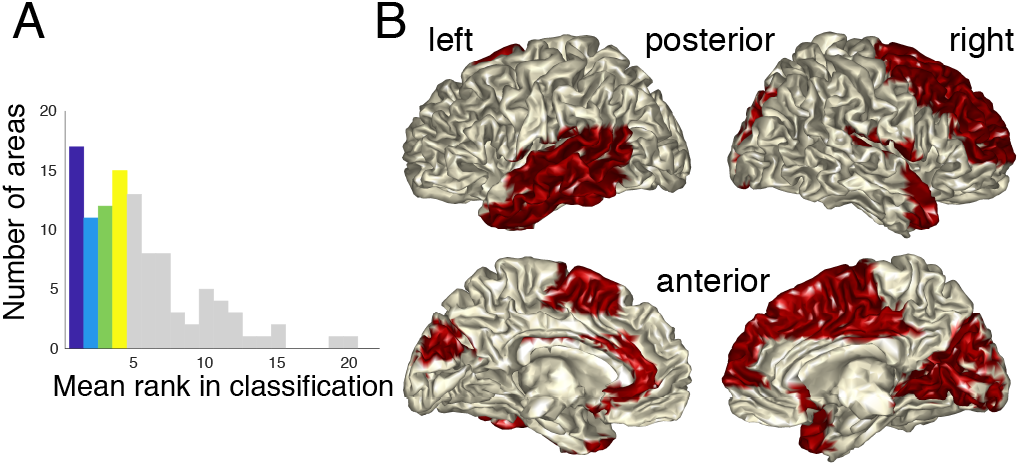
Cross-group comparison. (A) Histogram of classification ranks. Bin width is 1. (B) The topographic distribution of significantly different classification ranks in the cross-group classification is highlighted in red, as tested by a permutation procedure.

In the cross-group analysis, 54.8% of 115 areas obtained classification ranks ranging from 1 to 4, while the automatic identification of the remaining regions was less precise (see Fig. 4A). Beyond these descriptive procedures, we statistically compared classification results in the cross-group condition as compared to the within S-BF group classification (see Fig. S2 for the distributions of all brain areas). As seen in Fig. 4B, cross-group classification ranks were significantly worse for several sensory as well as for right frontal areas, and an extended network of left temporal brain regions. This suggests that spectral profiles in sensory (e.g., right calcarine, right Heschl’s gyrus, left superior temporal gyrus) and right frontal (e.g., right superior frontal gyrus) brain regions were different in the CB group as compared to the sighted, while no differences were found for other brain areas (see Table 1 for all ROIs showing significantly worse classification compared to the null-distribution). A control analysis was performed, to assure that group-differences were not introduced by the normalization procedure (i.e. reflecting differences in baseline power; Fig. S5). In the following, we focus on the interpretation of findings at ROIs that showed significant group differences in both cross-group classification analyses (i.e., with and without normalization procedure). These areas include: right Heschl’s gyrus, Calcarine gyrus, Cuneus, superior medial frontal gyrus, middle cingulate cortex and left middle temporal pole and anterior cingulate cortex (highlighted in Table 1).

**Table 1.** Table of all brain areas (out of 115) with significant classification differences. Brain areas where classification ranks were significantly different between the congenitally blind and the sighted (blindfolded) are listed. The areas highlighted in bold showed significant group differences in the cross-group classification analyses (i.e., with normalization procedure) and in the control analysis (i.e., without normalization procedure; see Fig. S5) and will be the focus of interpretation.

| Coarse region | Hemisphere |  |
| --- | --- | --- |
|  | Left | Right |
| Frontal |  | Superior frontal gyrus |
|  |  | Middle frontal gyrus<br>**Superior medial frontal**<br>Rolandic Operculum |
| **Visual** | Cuneus | **Calcarine gyrus**<br>**Cuneus**<br>Superior occipital gyrus<br>Lingual gyrus |
| **Temporal** | Superior temporal gyrus<br>Middle temporal gyrus<br>**Middle temporal pole**<br>Inferior temporal gyrus | **Heschl's gyrus**<br>Middle temporal pole<br>Superior temporal pole |
| **Sensorimotor** | Supplementary Motor Area | Supplementary Motor Area |
| **Non-cortical** | Olfactory<br>**Anterior cingulate cortex**<br>Cerebellum lobule VI<br>Cerebellar vermis 3<br>Cerebellar vermis 9 | Olfactory<br>**Middle cingulate cortex**<br>Cerebellum lobule X<br>Cerebellar vermis 3<br>Cerebellar vermis 9 |

Interestingly, the brain areas identified as showing group differences in the spectral profiles in the cross-group classification were characterized by clusters comprising peaks with increased power at higher frequencies in the CB as compared to the S-BF participants (for an exemplary selection of brain areas with significant effects, see Fig. 5A; spectra of all brain areas are shown in Fig. S1). This pattern of results was observed for the auditory (with more power in the alpha and beta band in the CB as compared to the sighted participants) and the right frontal areas (more power in the theta band). In visual brain areas, power peaks were reduced in the alpha band for the CB as compared to the S-BF participants, in contrast, power was increased in the low-gamma band. Post-hoc permutation tests were performed to confirm these observations. To test differences in spectral signatures between the S-BF and CB, the raw Fourier spectra (i.e., without applying the spectral clustering) were extracted und subjected to permutation statistics. Participants’ group assignment (S-BF vs. CB) was permuted randomly (1000 permutations) (Q = 0.05; false discovery rate [FDR] corrected *p*-value = .033; *p*-values < .033) (see Fig. 5B and Fig. S3). Although the differences in low-gamma power in the calcarine between the CB and the S-BF were not significant in the post-hoc tests, see Fig. S6.

**Figure 5.**
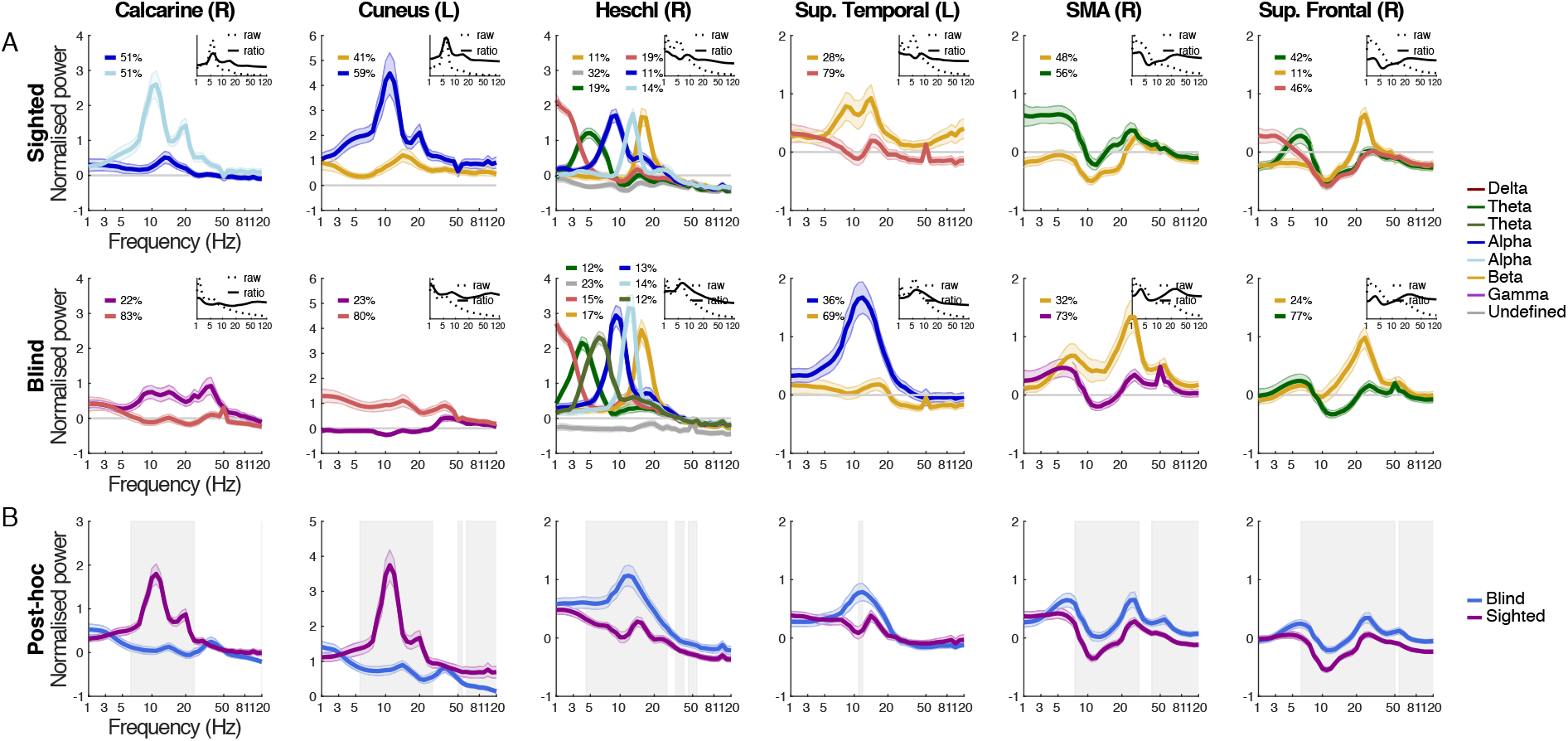
Comparison of spectral profiles of sighted and blind participants. (A) A selection of the spectral profiles of the brain areas (columns) that showed significantly worse cross-group classification is displayed, separately for the S-BF and the CB (rows). Clusters are color-coded according to the frequency of the cluster amplitude peak (legend on the right). The insets display the normalized mean power spectra (solid lines) and the unnormalized mean power spectra (dashed lines) (i.e. both without applying the spectral clustering). (B) Post-hoc analyses of the spectral differences are displayed for the selection of brain regions (columns). Spectra represent normalized mean power spectra (i.e. without applying the spectral clustering) at each ROI. Frequencies where power differences were obtained between the groups are indicated by grey boxes (permutation test: Q = 0.05; FDR corrected p-value = .033; p-values < .033). The groups are color-coded (legend on the right). In all panels: Shaded lines indicate standard error of the mean.

### Spectral Changes Correlate with Structural Group Differences

In order to test the relation of occipital spectral differences between the groups observed in the classifier analyses and their relation to brain structure, we performed a diffusion-tensor imaging (DTI) data analysis for a subsample of participants. The TBSS analysis revealed significantly higher Radial Diffusivity (RD) values in a bi-hemispheric spatial cluster in posterior parts of the brain (i.e., a cluster of voxels; displayed in Fig. S4 B) for the CB as compared to the sighted participants (*N*_*sighted*_ = 12, *N*_*CB*_ = 16; family-wise error (FWE)-corrected at the peak voxel, two-sided p = 0.05; see Fig. S4B), indicating reduced white matter structural integrity in the CB group. We used a probabilistic atlas of white matter pathways in MNI space (Thiebaut de Schotten et al. 2011) to evaluate the overlap of the spatial cluster with known white matter tracts. With the probabilistic atlas thresholded at 0.95, the TBSS cluster presents a significant overlap with the posterior corpus callosum, the posterior inferior longitudinal fasciculus (bilaterally), the posterior inferior fronto-occipital fasciculus (bilaterally), and the optic radiations (also bilaterally). This means that there is a 95% chance that the reduced white matter properties identified in the CB group by the TBSS analysis primarily affect these white matter tracts. Finally, RD values belonging to this spatial cluster were extracted and correlated with the power of individual cluster peaks from the spectral profile of occipital brain areas (Spearman correlations, FDR correction, Q = 0.05, corrected *p*-value = .0065, Fig. S4C). RD correlated negatively with the alpha power in right (rho = –0.55, *p* = .005) and left (rho = –0.81, p > 0.001) calcarine, as well as left cuneus (rho = –0.68, *p* > .001), indicating that reduced white matter properties in occipital areas were accompanied by reduced neuronal alpha power. Alpha and gamma power of the remaining occipital regions did not correlate with the RD values (all *ps* > .0065; cf. Table S1).

## Discussion

Intrinsic brain rhythms likely are involved in various sensory, motor and higher-cognitive processes. In congenitally blind individuals, behavioral performance in tasks involving the intact senses has often been found to be enhanced (Lessard et al. 1998; Roeder et al. 2000; Roeder et al. 2003; Gougoux et al. 2004; Hoetting et al. 2004; Roeder et al. 2004; Gougoux et al. 2005; Hertrich et al. 2009; Dietrich et al. 2013), and structural and functional reorganization has been observed in brain areas associated with the intact sensory systems (Pascual-Leone and Torres 1993; Sterr et al. 1998a; Sterr et al. 1998b; Roeder et al. 1999; Elbert et al. 2002; Stevens and Weaver 2005; Gougoux et al. 2009; Schepers et al. 2012; Hertrich et al. 2013; Watkins et al. 2013), as well as in brain regions predominantly associated with the visual system (Elbert et al. 2002; Burton 2003; Roeder and Neville 2003; Noppeney et al. 2005; Pascual-Leone et al. 2005; Noppeney 2007; Bedny et al. 2011; Voss and Zatorre 2012). Whether and how these behavioral and neuronal adaptations affect brain rhythms outside of occipital brain regions had remained unclear to date. Here, we identified spectral alterations in congenitally blind individuals aiming at advancing a mechanistic understanding of adaptations to blindness. Furthermore, our findings add to the ongoing debate regarding the functional relevance of brain rhythms. Confirming our first two hypotheses, spectral clustering and classification procedures performed exceptionally well for all groups (97–100% of areas were classified correctly in each group). This highlights consistent brain area-specific spectral properties across individuals within the sighted group and, as demonstrated for the first time, within the congenitally blind group. Crucially, cross-classification showed that visual deprivation co-occurred with changes in the spectral profiles especially of sensory (auditory and visual), right-frontal, left middle temporal pole and bilateral parts of the cingulate cortex. This was reflected in increased power in the theta-to-beta frequency bands in the right primary auditory cortex and right-frontal brain regions. In visual cortex, the sighted showed the alpha rhythm typically observed during rest, whereas in the congenitally blind individuals, clusters with expected decreased alpha power were accompanied by a gamma (~40 Hz) peak, suggesting an altered inhibitory-excitatory balance in the blind. Additionally, a correlation analysis of individual spectral clusters in occipital cortex and microstructural white matter properties revealed a positive correlation between alpha (but not gamma) power and white matter integrity in an extended bi-hemispheric spatial cluster in posterior parts of the brain.

### Robust Classification of Brain Areas Based on Spectral Profiles

Spectral profiles measured during resting state have revealed intrinsic brain rhythms that are thought to be crucially involved in task-related processing (Smith et al. 2009; Hipp et al. 2012; Engel et al. 2013; Raichle 2015; Keitel and Gross 2016; Sormaz et al. 2018). In the present study, we replicated the observation that brain areas expressed characteristic spectral profiles (Keitel and Gross 2016). Spectral clustering and automatic classification revealed spectral profiles, classification ranks, and distributions of classification ranks across the cortex in the sighted (S-EO) group similar to the ones first reported by Keitel and Gross (2016). Spectral profiles, for example of occipital regions, showed the typically observed alpha rhythm peaking at ~10 Hz, which has repeatedly been found to be involved in attentional inhibition (Klimesch et al. 2007; Buffalo et al. 2011; Jensen et al. 2012). Spectral peaks in the beta band (~20 Hz) were prominent across frontal and central brain areas, resembling previously reported natural frequencies of these brain areas (Rosanova et al. 2009; Ferrarelli et al. 2012; Keitel and Gross 2016), which may play a role in temporal motor processing (Fujioka et al. 2012; Arnal et al. 2015; Morillon and Baillet 2017).

While the spectral profiles of most brain areas resembled those reported by Keitel and Gross (2016), for some brain areas the spectral profiles differed (see Fig. S1). In our study, deep subcortical brain areas (in contrast to what has been reported by Keitel and Gross 2016) were not classified well (Fig. S1). A possible reason for this is that the slightly shorter segment duration used in the present study, which reduced the frequency resolution at lower frequencies, affected the detection of low-frequency clusters. Alternately, a lower signal-to-noise ratio in deeper brain areas in our data as compared to Keitel and Gross (2016) might be related to the employment of different MEG systems. Thus, possibly the recording system used and/or the sample of participants tested potentially influenced the specific profiles of some brain areas more than others. A test on a large dataset across different recording sites (i.e., several hundreds of recordings) will be necessary to clarify which spectral modes generalize across individuals of a larger population. Importantly, within our sample, the spectral profiles were consistent across individuals (i.e., only group clusters were reported in which at least ~70% of participants, and on average ~97% for the S-EO and ~94% for the S-BF group, contributed to each of the group-level spectral clusters). In summary, our results highlight the robustness of brain area-specific spectral profiles, suggesting that spectral profiles are characteristic properties reflecting the intrinsic brain rhythms of cortical regions.

On the premise that changes in spectral profiles are functionally related to adaptive plasticity in congenitally blind individuals, they are expected to be consistent (and not random) within a group of congenitally blind individuals. A novel finding of our study which is in accord with this hypothesis is that in the congenitally blind group, as with the sighted group, brain regions were identified reliably based on their spectral clusters, suggesting spectral consistencies across individuals (Fig. 2B, right; i.e., only group clusters were reported in which at least ~69% of participants and on average ~95 % contributed to each spectral cluster). This result suggests that adaptation of the cortex to visual deprivation leads to specifically and homogenously altered spectral fingerprints in the congenitally blind individuals.

### Selective Spectral Plasticity Across the Brain

The main goal of our study was a comprehensive (in space and frequency), data-driven model of spectral fingerprints in the congenitally blind as compared to sighted individuals in order to provide insights into the neuronal mechanisms underlying plasticity in congenital blindness. In the cross-group classification, brain areas of individual congenitally blind participants were classified based on the group-level spectral clusters of the sighted (S-BF). In order to isolate visual deprivation-related effects, the participant groups were well matched in our study (cf. methods section). While in the cross-group classification the classification for the majority of the brain areas was relatively good (i.e., low ranks; Fig. 4A), spectra related to right auditory, occipital and right frontal regions were classified significantly worse as compared to the within-sighted classification (Table 1; Fig. 4B). These findings suggest that spectral properties may have been systematically altered by visual deprivation-related plasticity for certain brain areas. Previously, a non-monotonic relationship between plasticity and stability across the cortex has been reported using fMRI (Haak and Beckmann 2019), whereas the plasticity observed in neuronal activity has been related to gene expressions (Ortiz-Terán et al. 2017). Specifically, plasticity likely decreases from early visual to mid-level cortex, but increases again further in the visual cortical hierarchy (Haak and Beckmann 2019).

### Spectral Plasticity in Sensory Areas

Altered spectral profiles in the congenitally blind reflect changes in intrinsic brain rhythms and might be linked to cognitive processes. Our findings highlight changes in spectral properties of the auditory and visual cortices due to visual deprivation-related neuroplasticity. The findings confirm previous reports demonstrating both cross-modal reorganization in visual cortex (Burton 2003; Pascual-Leone et al. 2005; Bedny et al. 2011; Voss and Zatorre 2012; Gudi-Mindermann et al. 2018; Rimmele et al. 2019) and intra-modal reorganization in auditory cortex (Roeder and Neville 2003) in blind humans. Importantly, our findings extend these results by providing evidence for genuine changes in the processing mode of these regions, as indicated by changes in the spectral characteristics, which contribute to a more mechanistic understanding.

Visual brain areas classified as spectrally different between the sighted and the congenitally blind individuals (in both the normalized and non-normalized control analysis) comprised right primary visual cortex (calcarine sulcus) and cuneus (Table 1). In these areas, we observed in the sighted two alpha-band clusters indicating distinct alpha brain rhythms. The posterior alpha rhythm, which is typically observed in visual areas during resting state, likely reflects a mechanism of rhythmic inhibition of task-irrelevant neuronal circuits (Haegens et al. 2011). In our data, we observed one cluster with a clear visual alpha peak at ~10 Hz and a smaller peak in the beta-band (~20 Hz), and a second alpha cluster characterized by a smaller amplitude (note that the two clusters are displayed by two separate lines in Fig. 5A), corroborating previous work suggesting distinct visual alpha rhythms (Barzegaran et al. 2017). In contrast, in the congenitally blind individuals these typical visual areas were characterized by a first cluster with a peak at higher frequencies (low gamma, ~40 Hz) and a strongly reduced alpha power peak, as well as a second cluster with low power in the alpha band (Fig. 5A, 5B). This observation is in line with previous findings reporting a reduced or entirely absent alpha rhythm in the visual system in blind individuals (Adrian and Matthews 1934; Noebels et al. 1978; Kriegseis et al. 2006; Hawellek et al. 2013). The reduced alpha brain rhythm is thought to reflect atrophy or reorganization of the cortico-thalamic and cortical pathways, where posterior alpha generators have been located (Lopes da Silva 1991; Lőrincz et al. 2009). It has been proposed that synchronized posterior gamma activity is controlled by alpha (de-)synchronization and that it indicates increased neuronal gain allowing feedforward processing of sensory information (van Kerkoerle et al. 2014; Michalareas et al. 2016). The gamma rhythm in sighted individuals is typically increased in the presence of a visual stimulus, but decreased or absent during rest. The gamma rhythm might be preserved in blind individuals, as it evidently emerges locally in the occipital cortex (Bastos et al. 2014; Marshall et al. 2018) being less impacted by atrophy of the thalamocortical pathway. The presence of gamma activity in blind individuals during rest possibly reflects the alteration of the alpha-gamma excitation-inhibition balance, i.e., the reduced alpha rhythm may result in a disinhibition of visual cortex increasing gamma band activity (Roeder et al.). Functionally, while the reduced alpha rhythm in the congenitally blind may reflect cortical and thalamo-cortical atrophy, the gamma-band rhythm might be related to the compensatory reorganization of visual cortex, i.e., the recruitment of visual cortex during the processing of non-visual tasks (Voss and Zatorre 2012; Striem-Amit et al. 2015; Bedny 2017; Voss 2019). This conjecture is inline with our correlation analysis of the spectral profiles with microstructural white matter properties (Fig. S4, Table S1), which showed that only the alpha cluster correlated with the deteriorated white matter properties.

Additionally, we found altered spectral profiles in right auditory cortex (Heschl’s gyrus; Table 1) with increases in power in specific frequency bands. In this area, we observed increased power in the theta-to-beta frequencies in blind as compared to sighted participants (Fig. 5A, 5B). Importantly, in a comparison of auditory cortex spectral profiles during rest and during speech comprehension, Keitel and Gross (2016) provided evidence for the functional relevance of intrinsic delta, theta and beta brain rhythms, as observed during rest, for speech processing. Speech tracking in the theta band (i.e., the phase-locking of auditory cortex oscillatory activity to speech acoustics) is tightly linked to the temporal segmentation of syllables, as well as to the prediction of upcoming stimuli (Giraud and Poeppel 2012; Gross et al. 2013; Haegens and Zion Golumbic 2018; Rimmele, Gross, et al. 2018; Rimmele, Morillon, et al. 2018). Moreover, these auditory cortex brain rhythms seem crucial for the segmentation of sound and music (Doelling and Poeppel 2015; Morillon and Baillet 2017). Interestingly, previous research has shown that congenitally blind individuals are able to comprehend ultra-fast speech at rates where speech comprehension fails in sighted individuals (i.e., speech comprehension in sighted individuals has been shown to fail at syllable rates faster than the theta range; Brungart et al. 2007; Ghitza and Greenberg 2009). Furthermore, this ability to process ultra-fast speech in the congenitally blind individuals has been linked to accelerated theta brain rhythms in right auditory cortex (Hertrich et al. 2013) (see also: Trouvain 2007; Hertrich et al. 2009; Dietrich et al. 2013; Van Ackeren et al. 2018). A possibility is that the altered spectral profiles in congenitally blind individuals reflect intrinsic brain rhythms that aid auditory temporal segmentation skills. Future research, however, is required to confirm this speculation by relating the changes in brain rhythms to behavioral performance.

### Spectral Plasticity Beyond Sensory Cortices

Beyond spectral reorganization in sensory cortices, our data suggest that also right-hemispheric frontal brain regions, bilateral parts of the cingulate cortex (left anterior, and right middle) and the left middle temporal pole undergo adaptation. Spectral clusters of right medial superior frontal gyrus and cingulate cortices were classified as different between the blind and the sighted groups, with increased power in the theta-band in the blind group. Medial superior frontal gyrus has been linked to various cognitive processes (Cohen et al. 2009; Cavanagh and Frank 2014; Chander et al. 2016; Töllner et al. 2017), as well as theta-band brain rhythms (Kubota et al. 2001; Onton et al. 2005; Cohen et al. 2008; Cohen 2011; Itthipuripat et al. 2013; Hsieh and Ranganath 2014; Töllner et al. 2017; Eschmann et al. 2018; Başar-Eroglu et al.). For instance, medial superior frontal gyrus has been implicated in cognitive control (Cavanagh and Frank 2014), cognitive control of timing (Lewis and Miall 2003), decision making (Schmidt et al. 2018), working memory (Itthipuripat et al. 2013; Hsieh and Ranganath 2014), temporal order processing (Chander et al. 2016), or language processing (Binder et al. 2009). Some of these processes are presumably altered in congenital blind individuals as compared to sighted individuals, such as language processing in a frontotemporal network (Roeder et al. 2002; Lane et al. 2017), and temporal order processing (Roeder et al. 2004; Stevens and Weaver 2005).

The cingulate cortex (among other processes) has been associated with emotion, cognitive control and pain processing (Shackman et al. 2011). The left temporal pole, part of the anterior temporal lobe, is thought to be involved in various processes including semantic (Binder et al. 1999; Visser and Lambon Ralph 2011) and conceptual (Baron and Osherson 2011) processing. For both the cingular (Ortiz-Terán et al. 2016) and anterior temporal (Striem-Amit et al. 2018) structures altered brain responses have been observed and proposed to be related to adaptive behavior in congenitally blind as compared to sighted individuals. However, as the functionality of these structures in congenital blindness, and particularly the role of endogenous brain rhythms, remain little understood, we refrain from further interpreting these findings.

Note, that for some brain areas we found differences in spectral profiles in the cross-group classification analysis (normalized data), however, not in the control analysis (on the non-normalized data; table 1; Fig. S5). We applied a previously established (Keitel & Gross, 2016) procedure. A disadvantage of the procedure is that it might introduce spurious activity due to group differences in the normalization spectrum, as the procedure normalizes the spectra across the brain. An advantage of the procedure is that it down-weights the neuronal activity (1/f power) that is predominant across the brain and not specific to a given brain area, which otherwise complicates the spectral analysis. Thus, without the normalization the noise-level of the data is increased due to individual differences in overall power across the spectrum and across the brain. Although these arguments justify our choice, as precautionary measure we only interpreted effects found in both analyses.

### Spectral profiles related to microstructural white matter properties

A relevant question is to what extent our findings of altered spectral profiles of occipital cortex in the congenitally blind are linked to structural differences. Our DTI analysis revealed compromised white matter integrity in congenitally blind individuals in visual association tracts comprising the ventral visual stream. These tracts included the bilateral inferior fronto-occipital fasciculus, connecting occipital and frontal brain areas and the bilateral inferior longitudinal fasciculus, connecting occipital and anterior temporal cortices, which have been conjectured to play a role in reading, writing and language semantics (Catani and Mesulam 2008); and the bilateral optic radiations, linking the visual thalamus to the primary visual cortex. In addition, white matter integrity was also compromised in the posterior corpus callosum, by which homologous visual cortices are interconnected (Restani and Caleo 2016). Reduced white matter integrity in early (Shimony et al. 2006; Park et al. 2007), late (Wang et al. 2013; Hofstetter et al. 2019), and congenitally blind individuals (Ptito et al. 2008; Wang et al. 2013; Aguirre et al. 2016; Reislev et al. 2016), particularly in visual pathways (e.g. optic radiations and the corpus callosum), is a well-documented finding. As myelination is proportional to the degree of neuronal activation during brain development (Demerens et al. 1996; Stevens et al. 2002; Fields 2004; Ishibashi et al. 2006; Gautier et al. 2015; Wake et al. 2015), the lack of visual processing during development in congenital blindness likely causes the reduced white matter density in visual pathways (Hill et al. 2014; Dietz et al. 2016; Restani and Caleo 2016; Anurova et al. 2019). Our findings showed that reduced spectral power in the visual alpha rhythm co-occurred with reduced white matter integrity in pathways that overlap with visual cortical pathways and thalamo-cortical projections (i.e., negative correlation between alpha power and RD; Fig. S4B), confirming previous accounts suggesting that reduced visual alpha power originates in structural impairment (Adrian and Matthews 1934; Noebels et al. 1978; Kriegseis et al. 2006; Hawellek et al. 2013).

### Concluding Remarks

The present study supports the findings of robust brain area-specific spectral profiles in the human brain. We provide, for the first time, a whole-brain model of spectral fingerprints in congenitally blind adults, and crucially, we were able to map spectral changes due to visual deprivation to specific—namely right visual, auditory, frontal, left anterior temporal, and bilateral cingulate—brain areas. We suggest that the power increases in the theta-to-beta frequency bands in auditory and frontal brain regions may reflect adaptive sensory or higher cognitive processing in blind individuals, while altered spectral profiles in visual brain regions (lower alpha and a gamma peak) may indicate a change in the excitation-inhibition balance. While these interpretations are tentative, the results pave the way for more targeted, task-based studies that aim to identify the link between altered spectral profiles and adaptive behavior.

## Acknowledgements

We wish to thank Laura Gwilliams, Federico Adolfi, and David Poeppel for their methodological and conceptual discussions and comments.

