## Supplement for "Data-Driven Classification of Spectral Profiles Reveals Brain Region-Specific Plasticity"

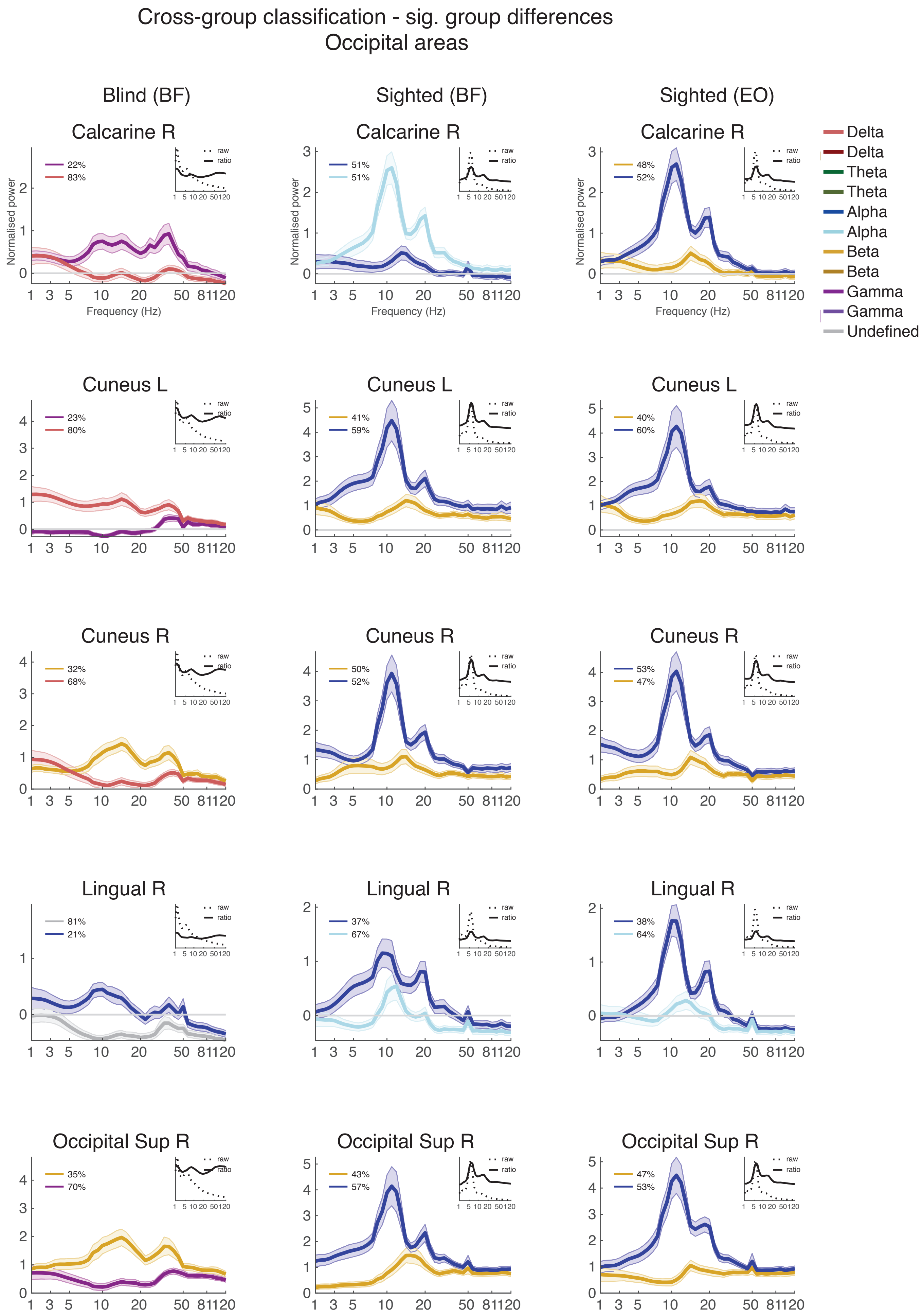
Cross-group classification - sig. group differences
Occipital areas
Blind (BF)
Sighted (BF)
Sighted (EO)
Calcarine R
Cuneus L
Cuneus R
Lingual R
Occipital Sup R
Normalised power
Frequency (Hz)
raw
ratio
22%
83%
51%
51%
48%
52%
23%
80%
41%
59%
40%
60%
32%
68%
50%
52%
53%
47%
81%
21%
37%
67%
38%
64%
35%
70%
43%
57%
47%
53%
Delta
Delta
Theta
Theta
Alpha
Alpha
Beta
Beta
Gamma
Gamma
Undefined

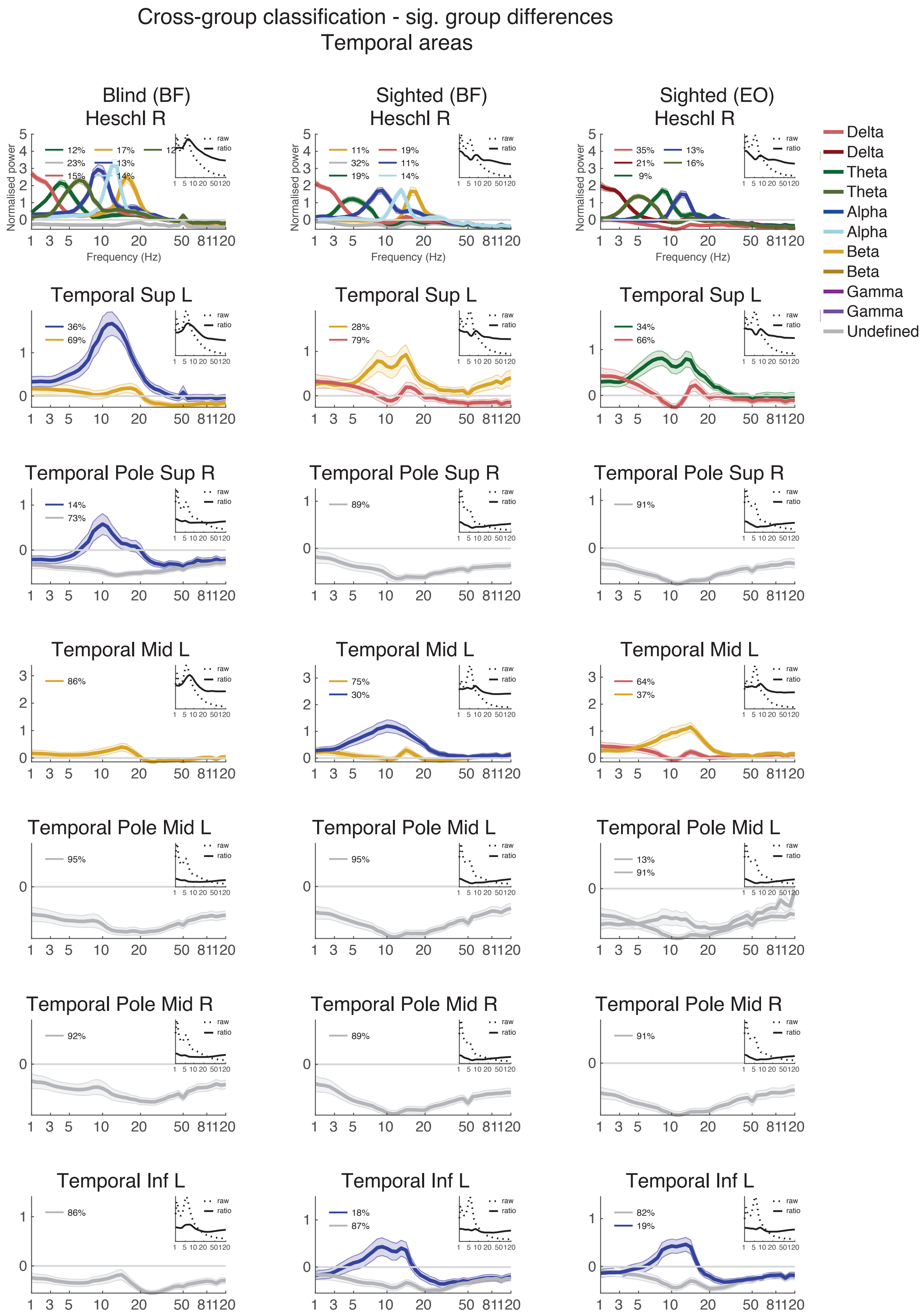

Cross-group classification - sig. group differences
Temporal areas
Blind (BF)
Sighted (BF)
Sighted (EO)
Heschl R
Temporal Sup L
Temporal Pole Sup R
Temporal Mid L
Temporal Pole Mid L
Temporal Pole Mid R
Temporal Inf L
Normalised power
Frequency (Hz)
raw
ratio
Delta
Delta
Theta
Theta
Alpha
Alpha
Beta
Beta
Gamma
Gamma
Undefined

### Cross-group classification - sig. group differences
### Frontal + non-cortical areas

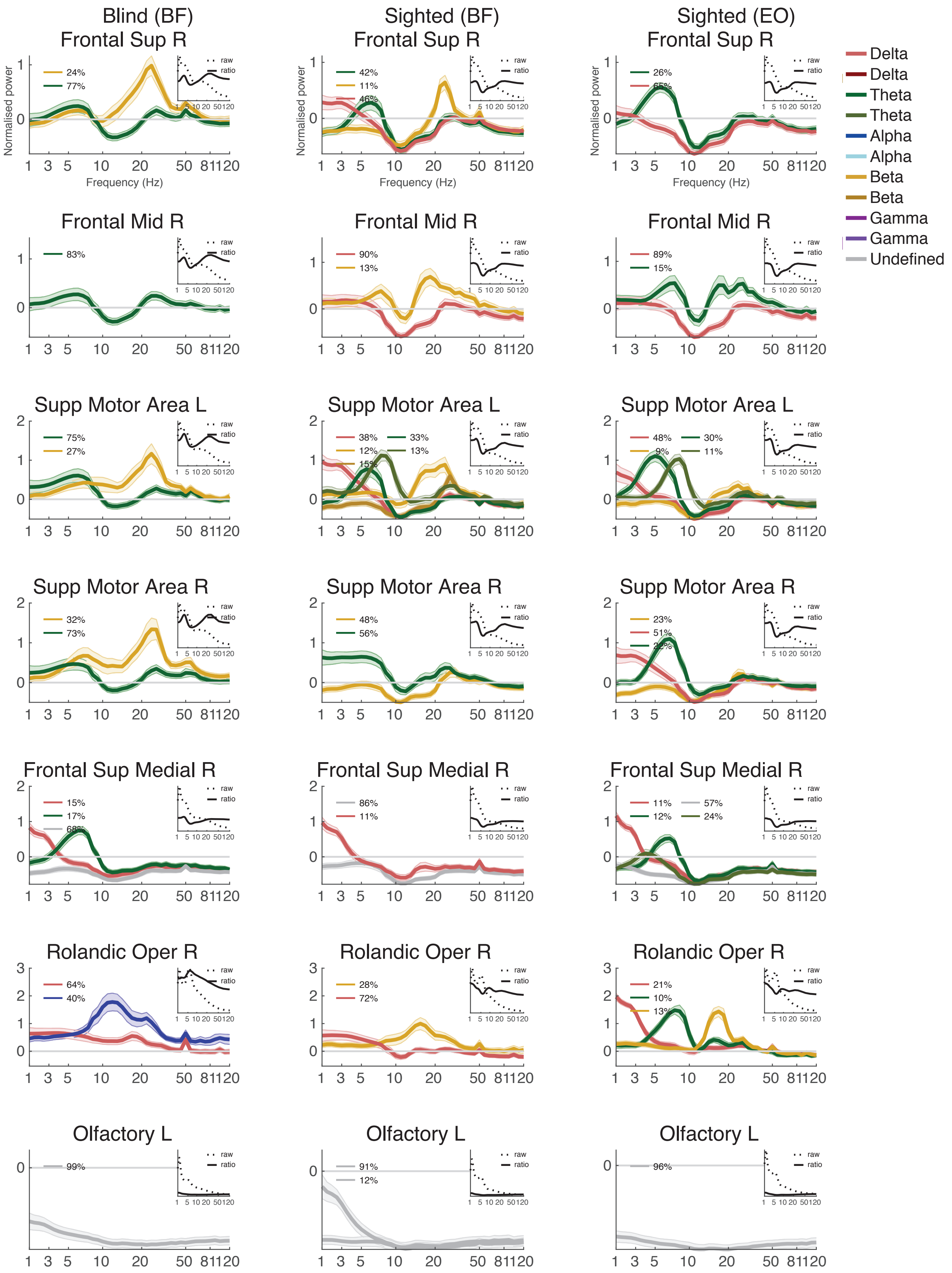

Cross-group classification - sig. group differences
Non-cortical areas

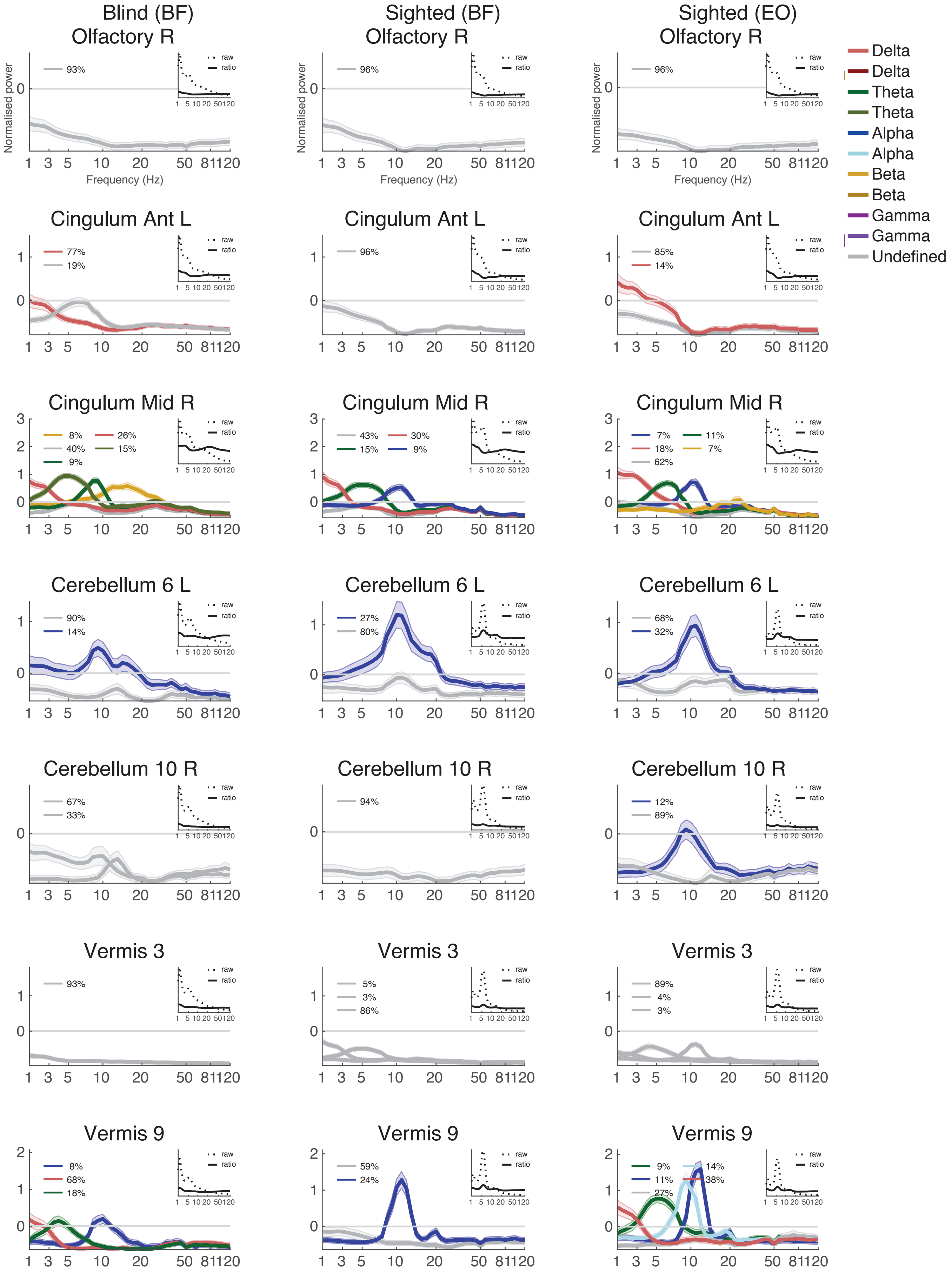

Cross-group classification - no sig. group differences
Occipital areas

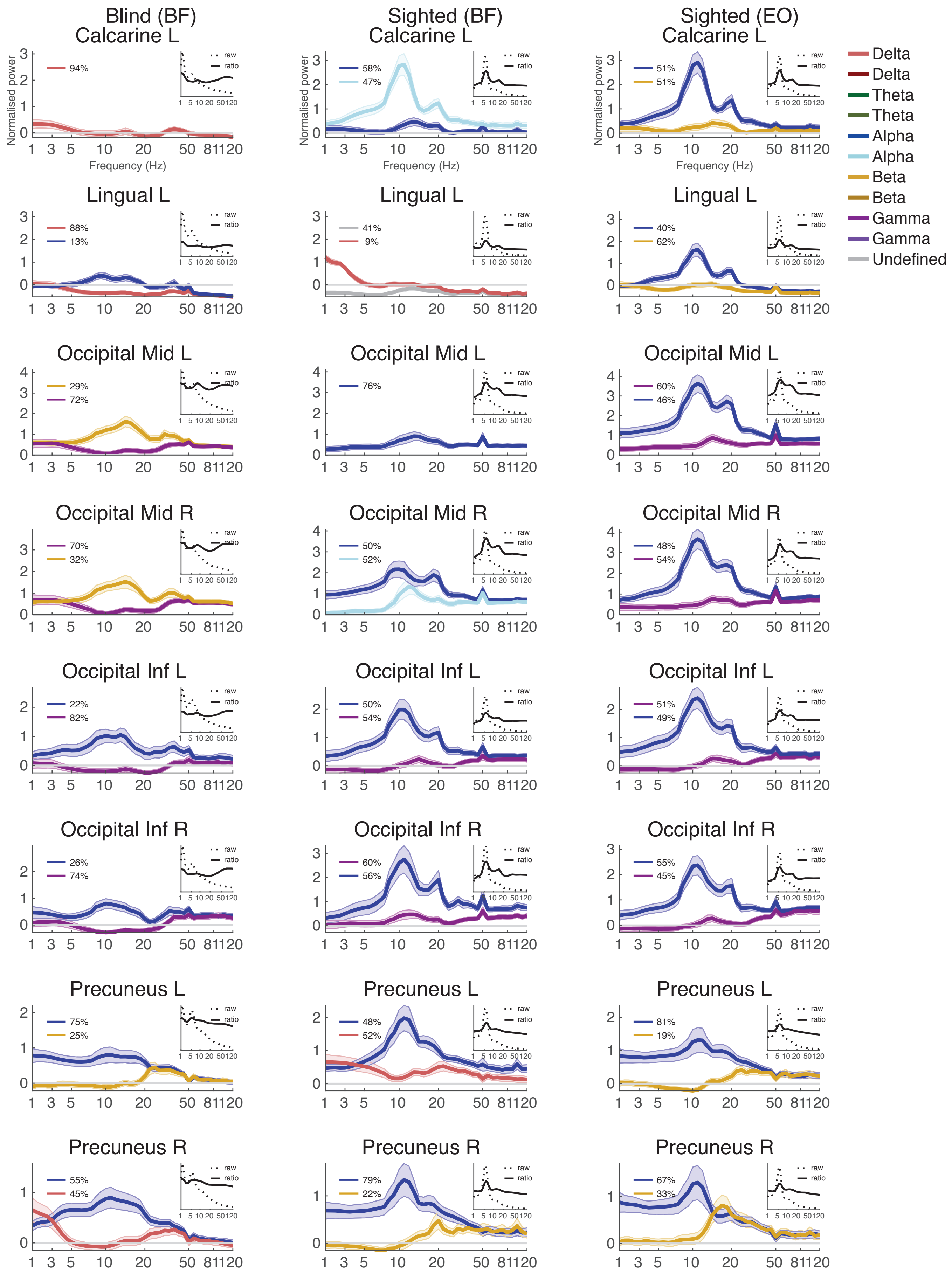

Cross-group classification - no sig. group differences
Temporal areas

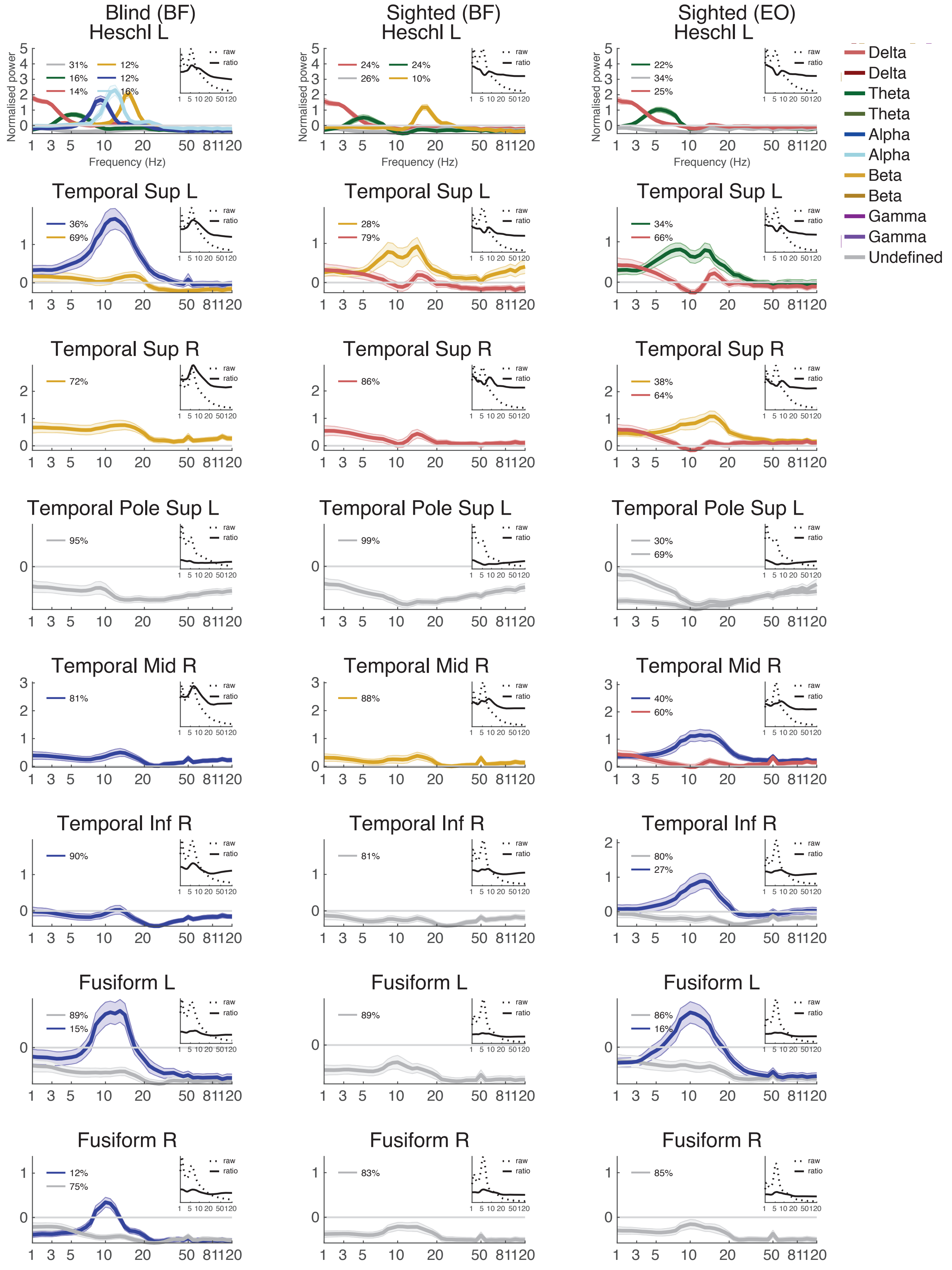

Cross-group classification - no sig. group differences
Central areas

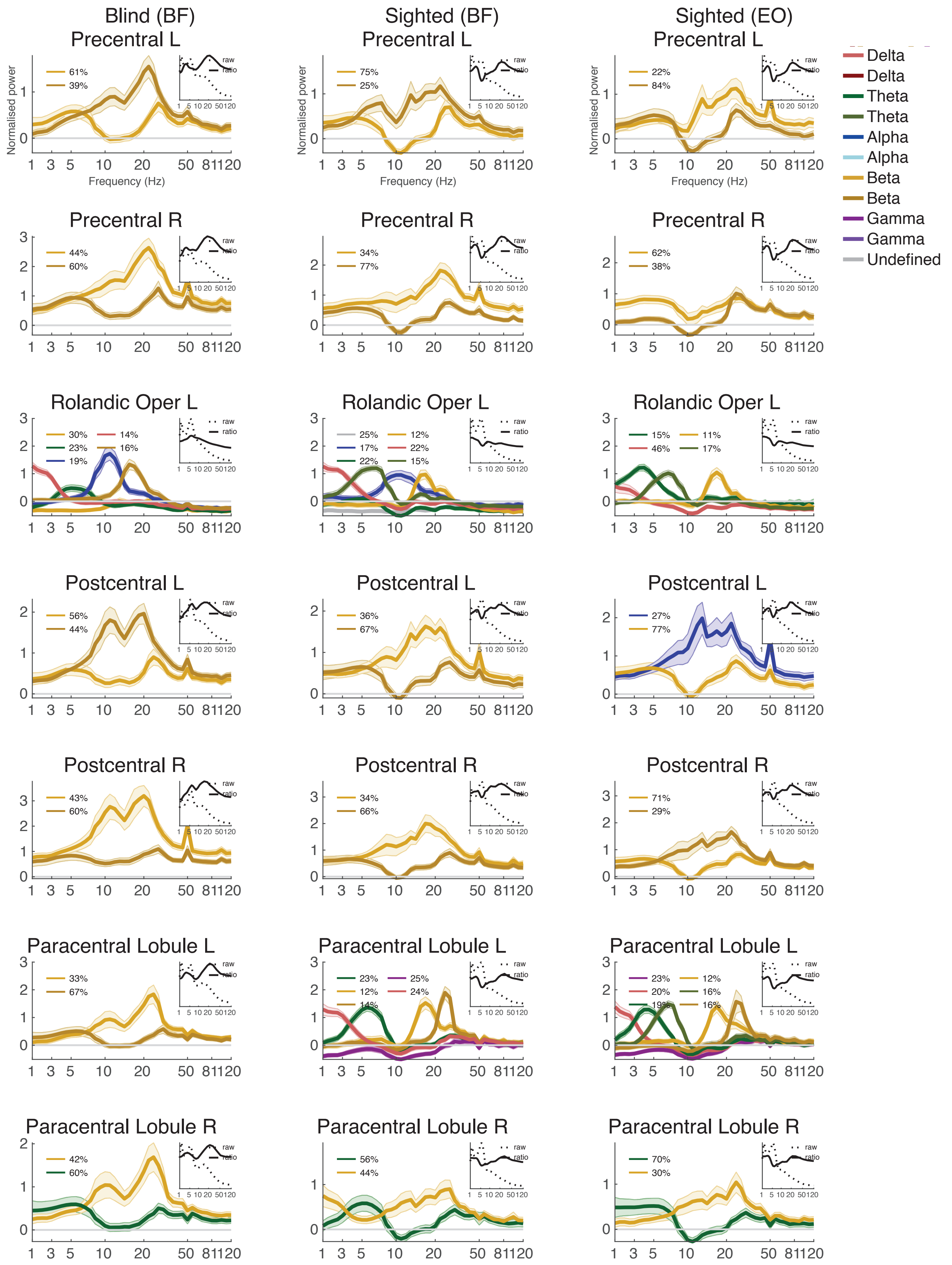

### Cross-group classification - no sig. group differences
### Central + parietal areas

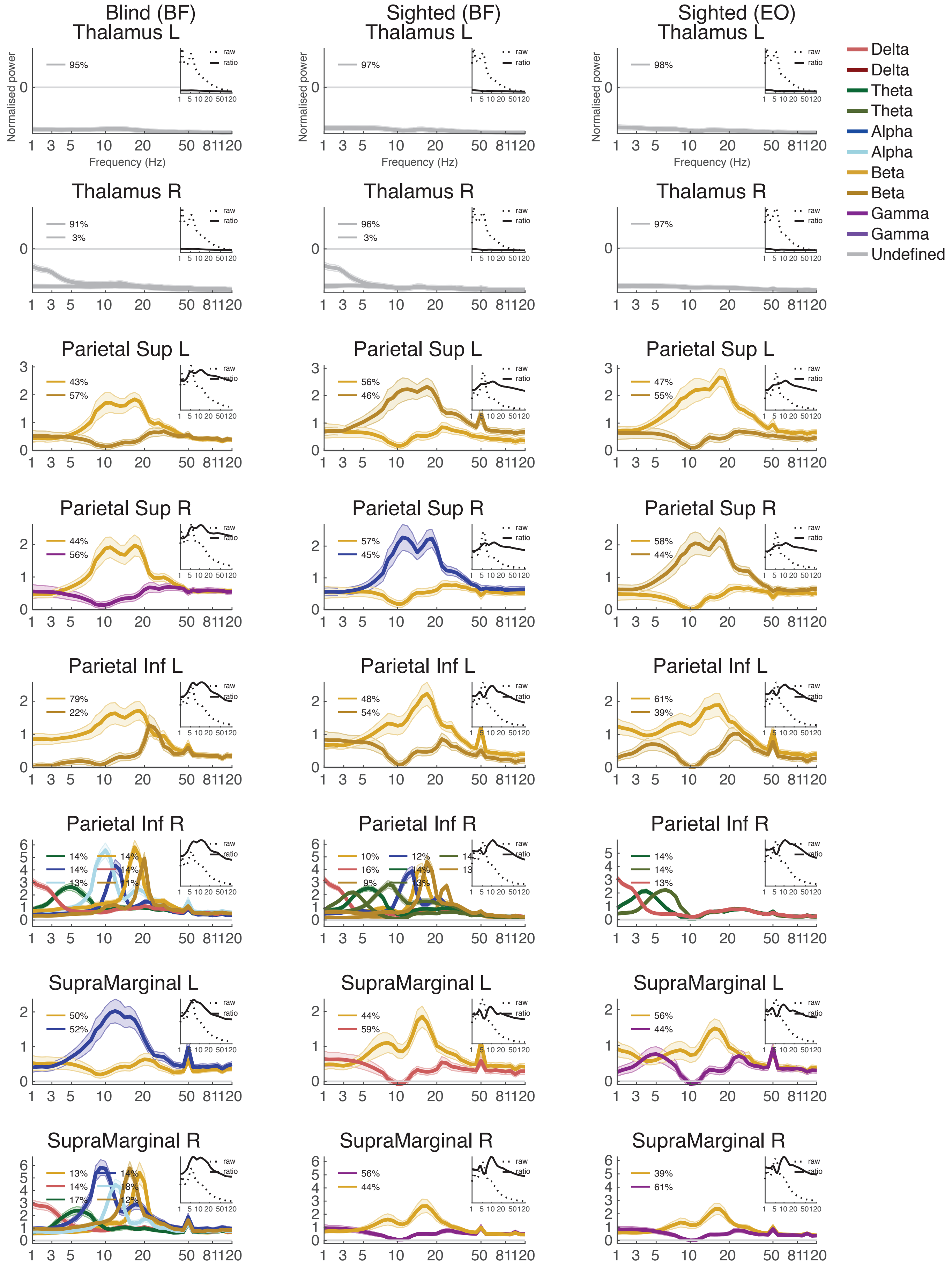

Cross-group classification - no sig. group differences
Parietal + frontal areas

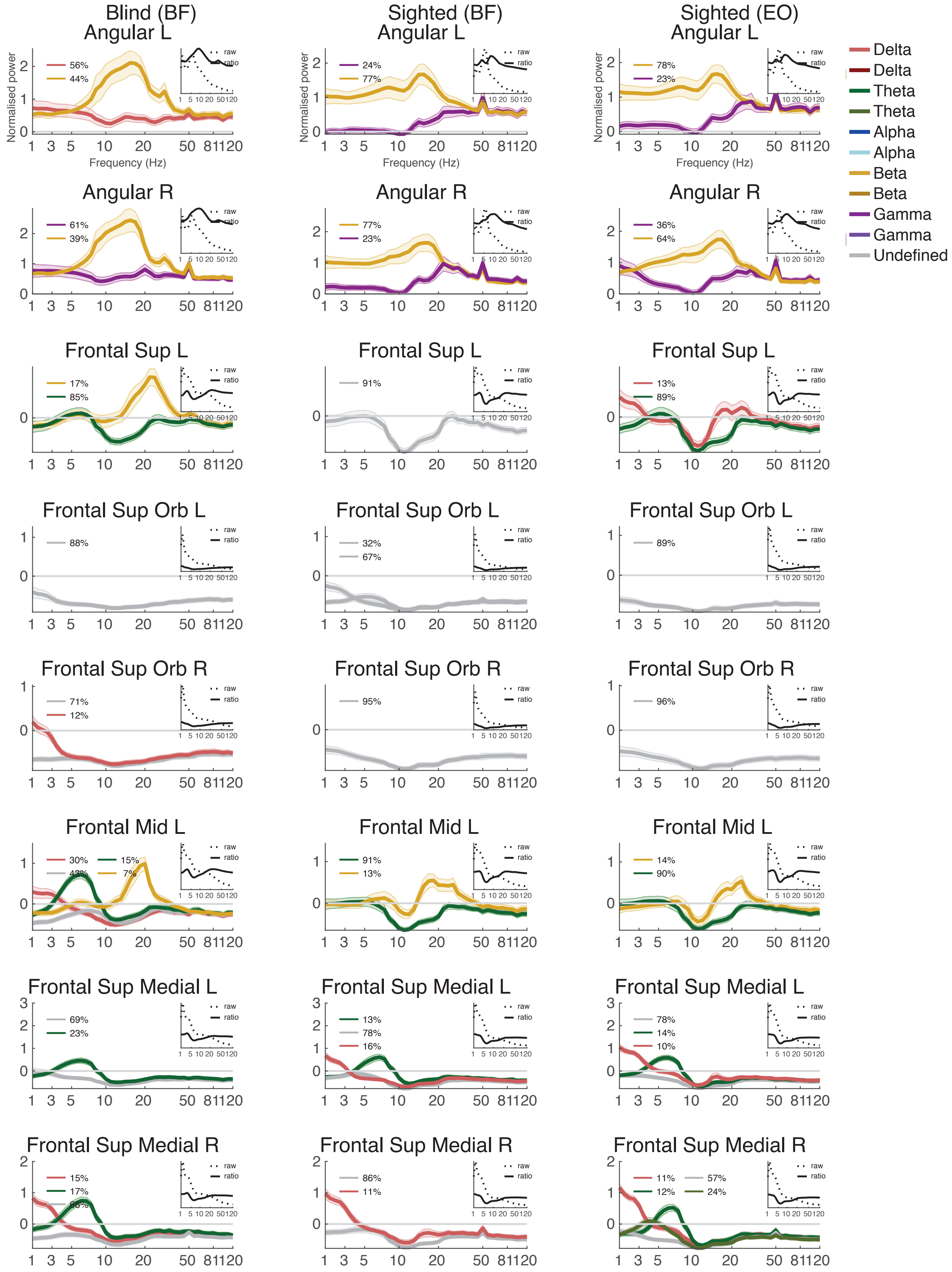

Cross-group classification - no sig. group differences
Frontal areas

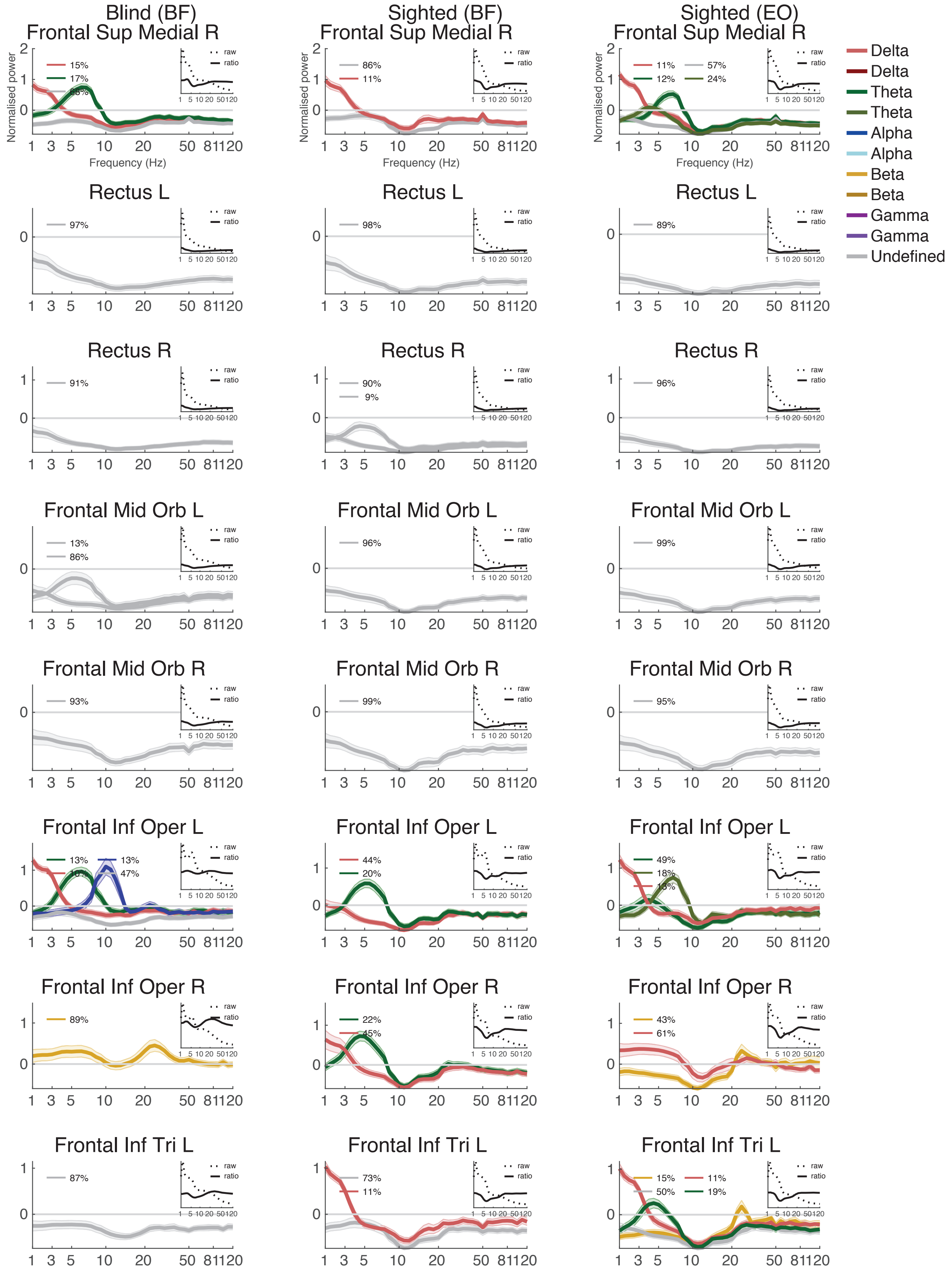

Cross-group classification - no sig. group differences
Frontal areas

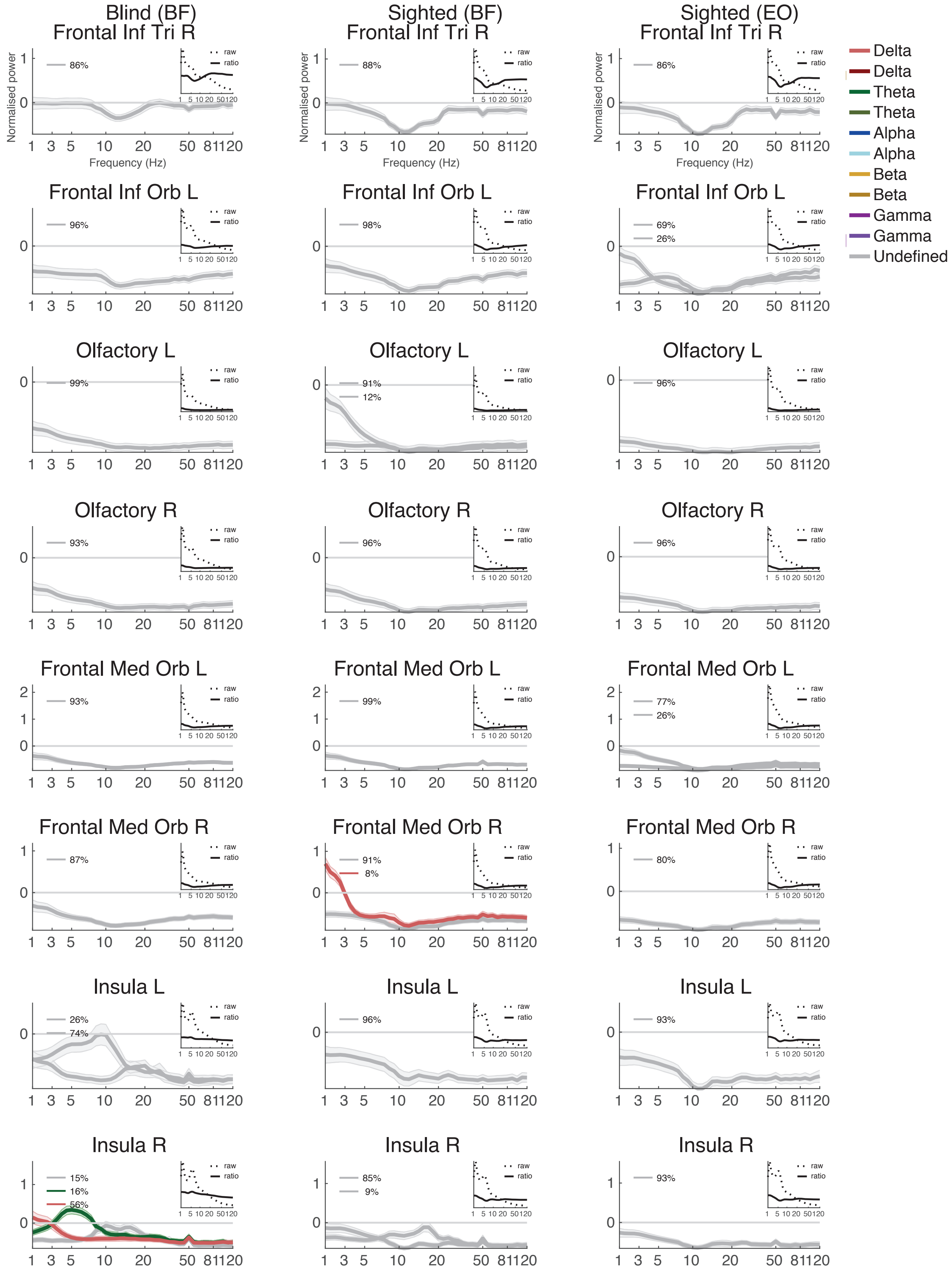

**Fig. S1. Spectral profiles of all brain areas**. Clustered spectral profiles shown separately for brain areas (rows) and experimental groups (columns). The x-axis illustrates frequency, scaled logarithmically from 1 to 120 Hz; the y-axis illustrates normalized power. Note that some clusters show spikes at 50 Hz and 100 Hz which are due to line noise. Each line corresponds to a cluster forming part of a spectral profile. The legends illustrate cluster durations, i.e., the percentage of single subject segments contributing to a group cluster, averaged across subjects. Clusters are color-coded according to the peak frequency (red: delta, green: theta, blue: alpha, yellow: beta, grey: no clearly-identifiable peak). Multiple clusters belonging to one frequency range are illustrated by different shades of the same color, i.e. dark and light blue. Shaded error bars represent the standard error of the mean across participants. Only group clusters to which the majority of subjects contributed are shown (i.e., CB: N=18; S-BF: N=17; S-EO: N=16). The insets display the normalized mean power spectra (solid lines) and the unnormalized mean power spectra (dashed lines) (i.e., both without applying the spectral clustering).

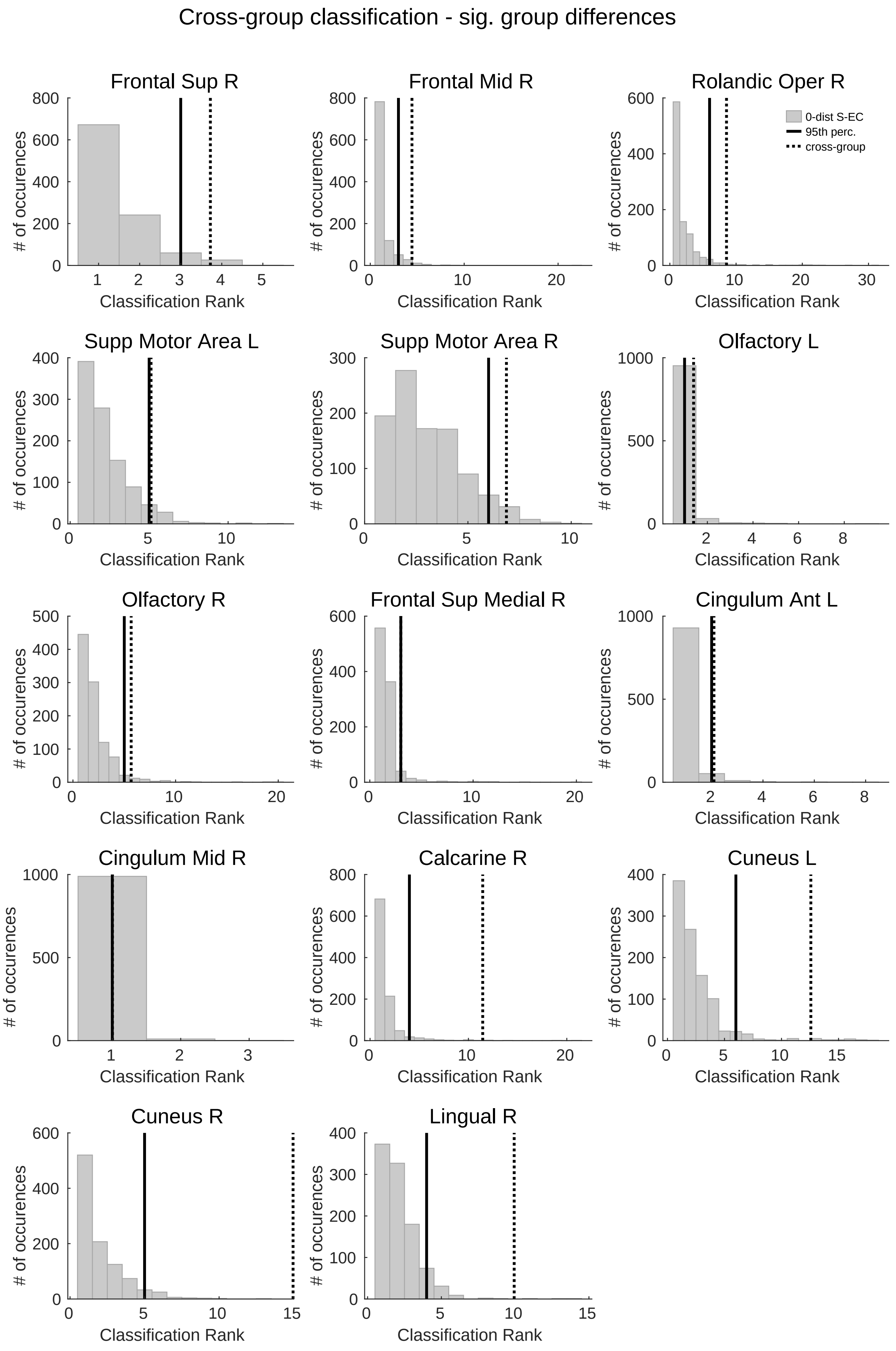
Cross-group classification - sig. group differences
Frontal Sup R
Frontal Mid R
Rolandic Oper R
0-dist S-EC
95th perc.
cross-group
Supp Motor Area L
Supp Motor Area R
Olfactory L
Olfactory R
Frontal Sup Medial R
Cingulum Ant L
Cingulum Mid R
Calcarine R
Cuneus L
Cuneus R
Lingual R
### of occurences
Classification Rank

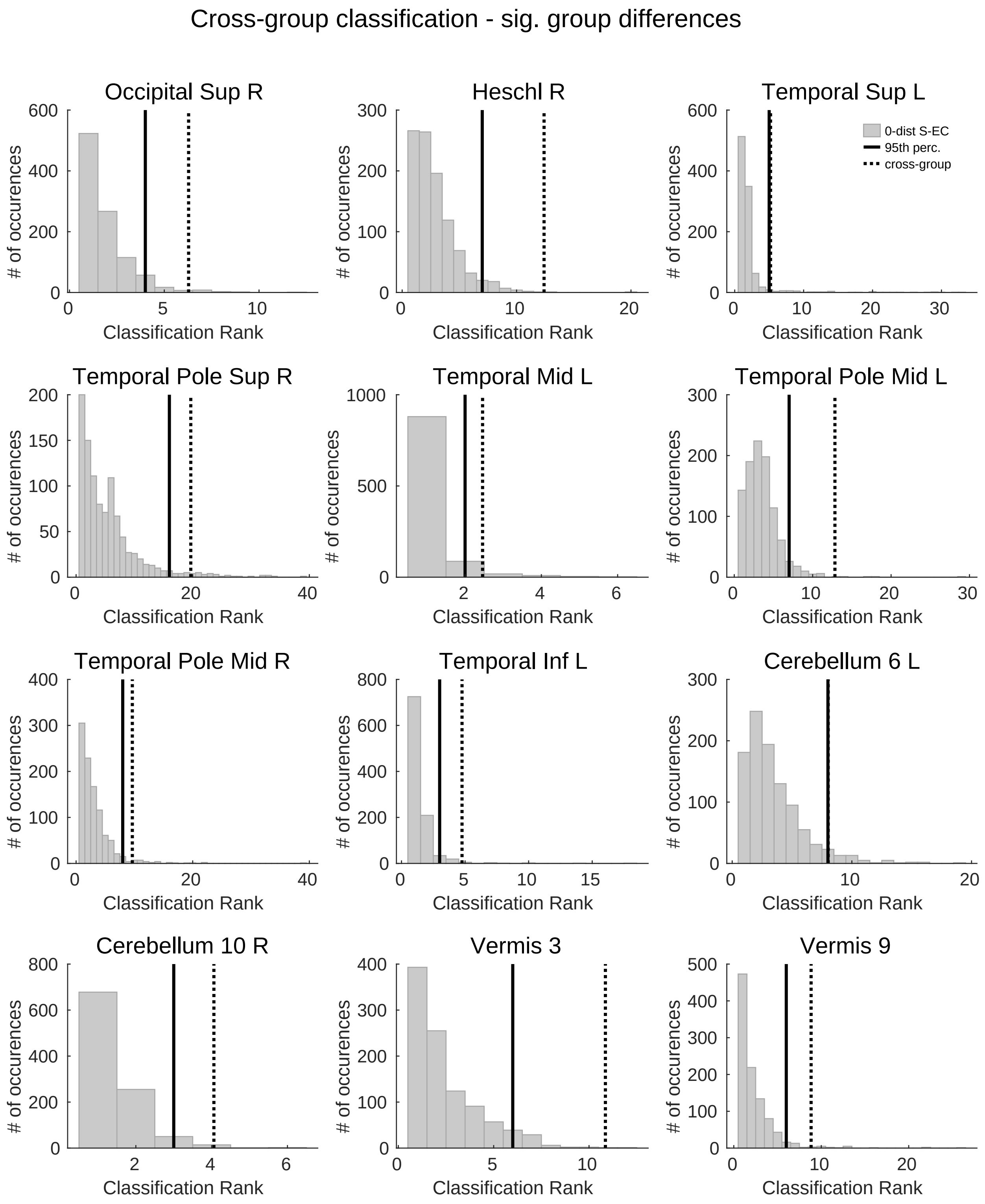

Cross-group classification - sig. group differences
Occipital Sup R
Heschl R
Temporal Sup L
0-dist S-EC
95th perc.
cross-group
Temporal Pole Sup R
Temporal Mid L
Temporal Pole Mid L
Temporal Pole Mid R
Temporal Inf L
Cerebellum 6 L
Cerebellum 10 R
Vermis 3
Vermis 9
### of occurences
Classification Rank

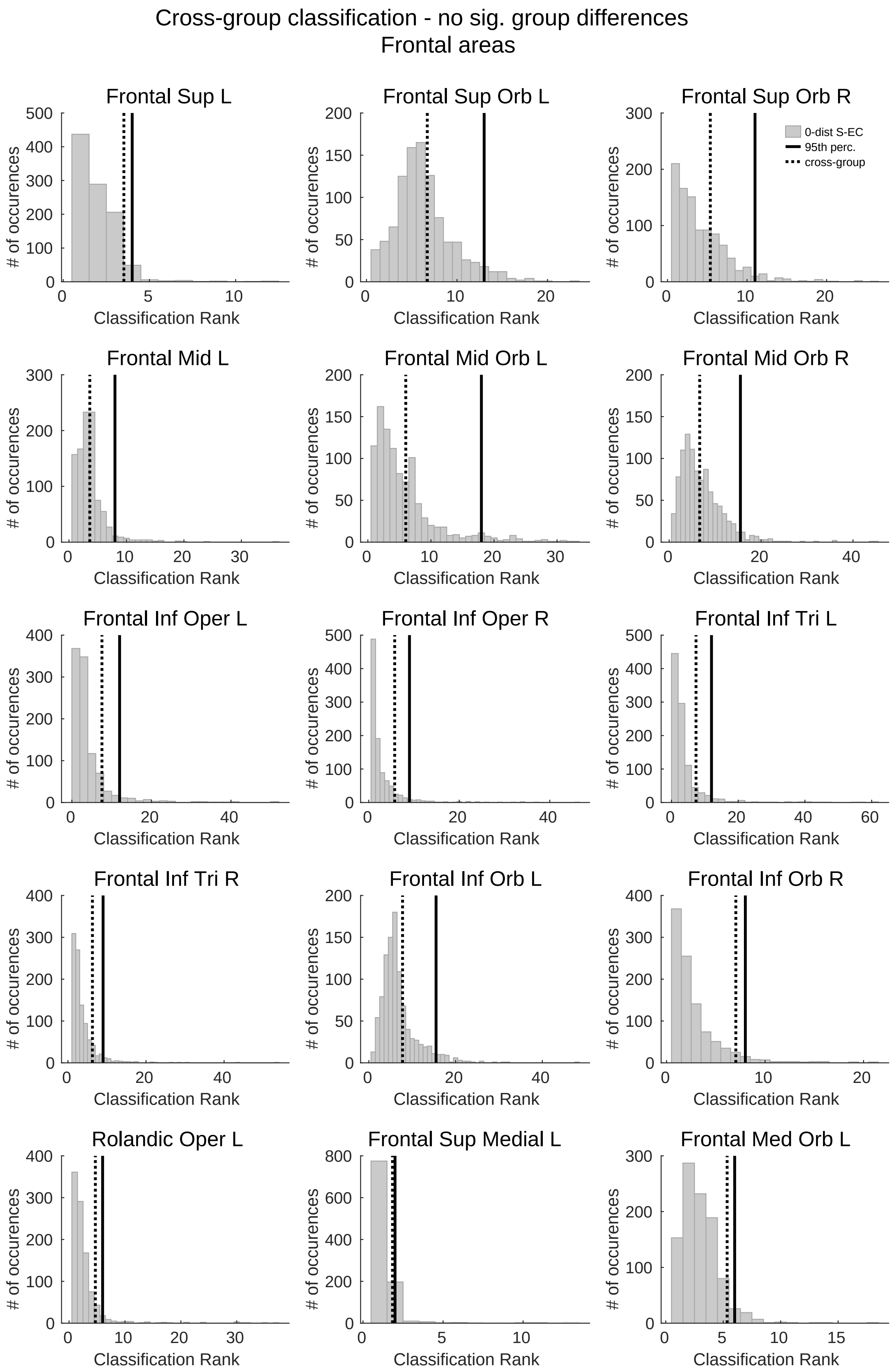

Cross-group classification - no sig. group differences
Frontal areas
Frontal Sup L
Frontal Sup Orb L
Frontal Sup Orb R
0-dist S-EC
95th perc.
cross-group
Frontal Mid L
Frontal Mid Orb L
Frontal Mid Orb R
Frontal Inf Oper L
Frontal Inf Oper R
Frontal Inf Tri L
Frontal Inf Tri R
Frontal Inf Orb L
Frontal Inf Orb R
Rolandic Oper L
Frontal Sup Medial L
Frontal Med Orb L
### of occurences
Classification Rank

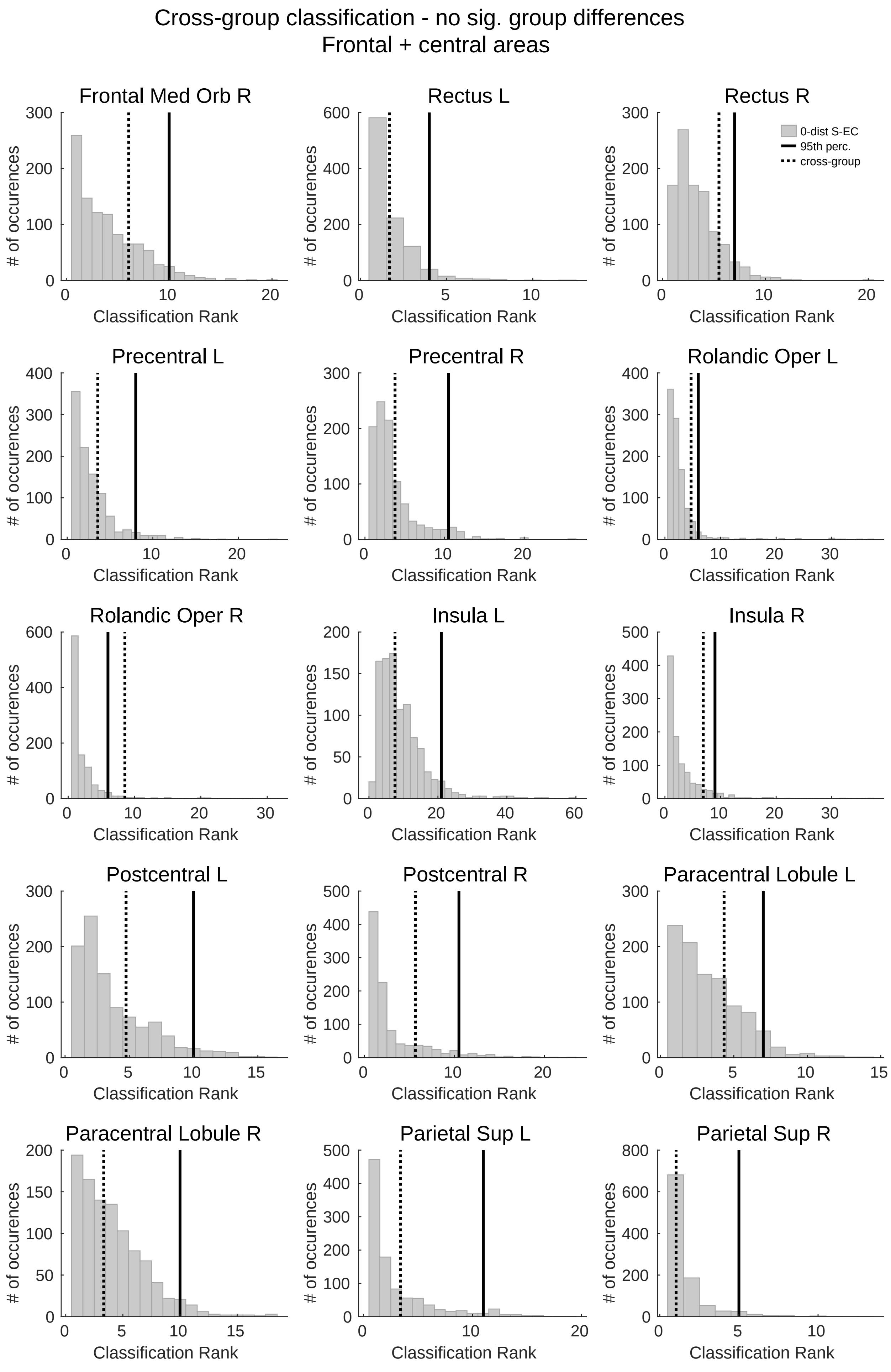
Cross-group classification - no sig. group differences
Frontal + central areas
Frontal Med Orb R
Rectus L
Rectus R
0-dist S-EC
95th perc.
cross-group
Precentral L
Precentral R
Rolandic Oper L
Rolandic Oper R
Insula L
Insula R
Postcentral L
Postcentral R
Paracentral Lobule L
Paracentral Lobule R
Parietal Sup L
Parietal Sup R
### of occurences
Classification Rank

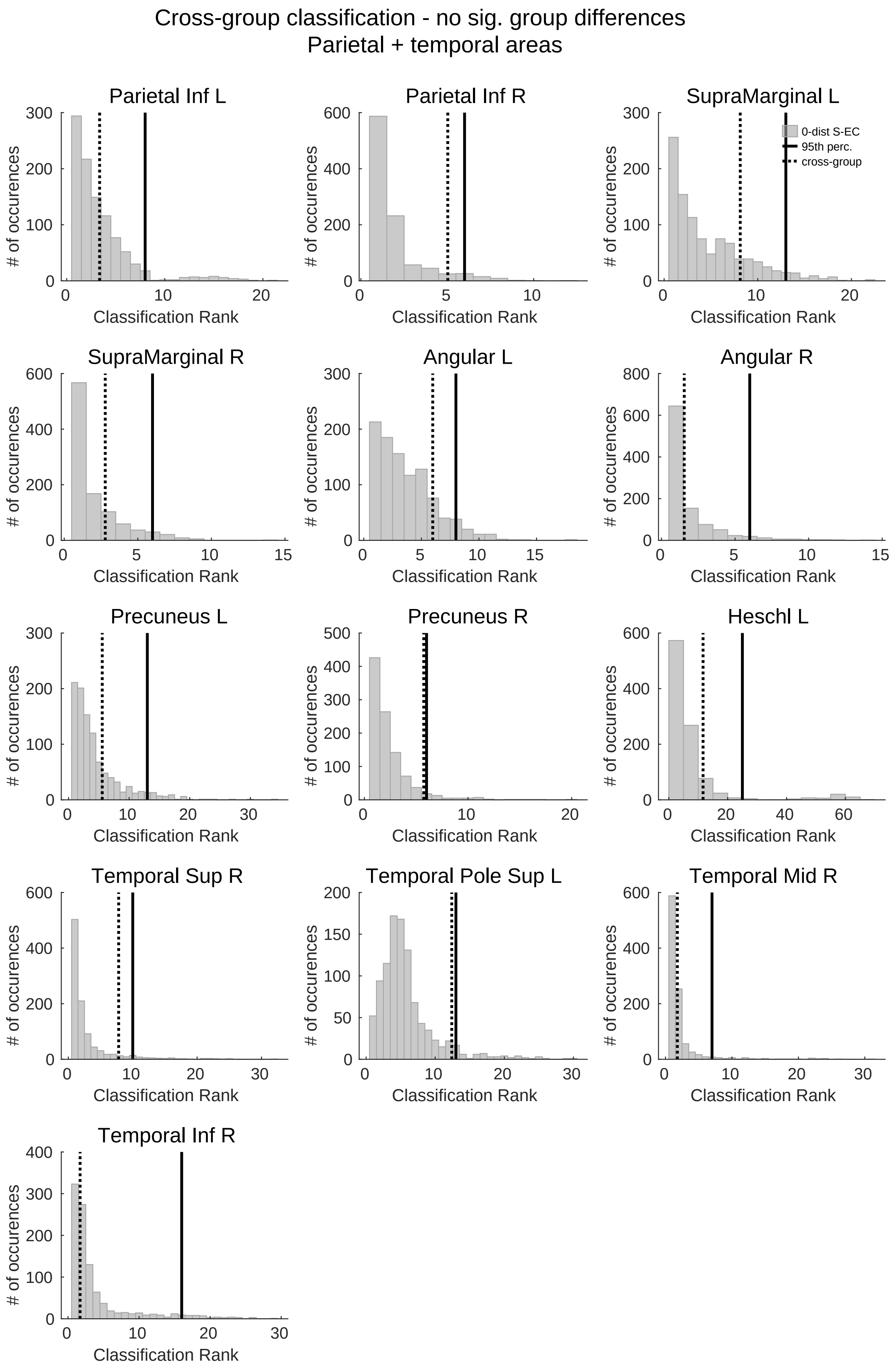
Cross-group classification - no sig. group differences
Parietal + temporal areas
Parietal Inf L
Parietal Inf R
SupraMarginal L
0-dist S-EC
95th perc.
cross-group
SupraMarginal R
Angular L
Angular R
Precuneus L
Precuneus R
Heschl L
Temporal Sup R
Temporal Pole Sup L
Temporal Mid R
Temporal Inf R
### of occurences
Classification Rank

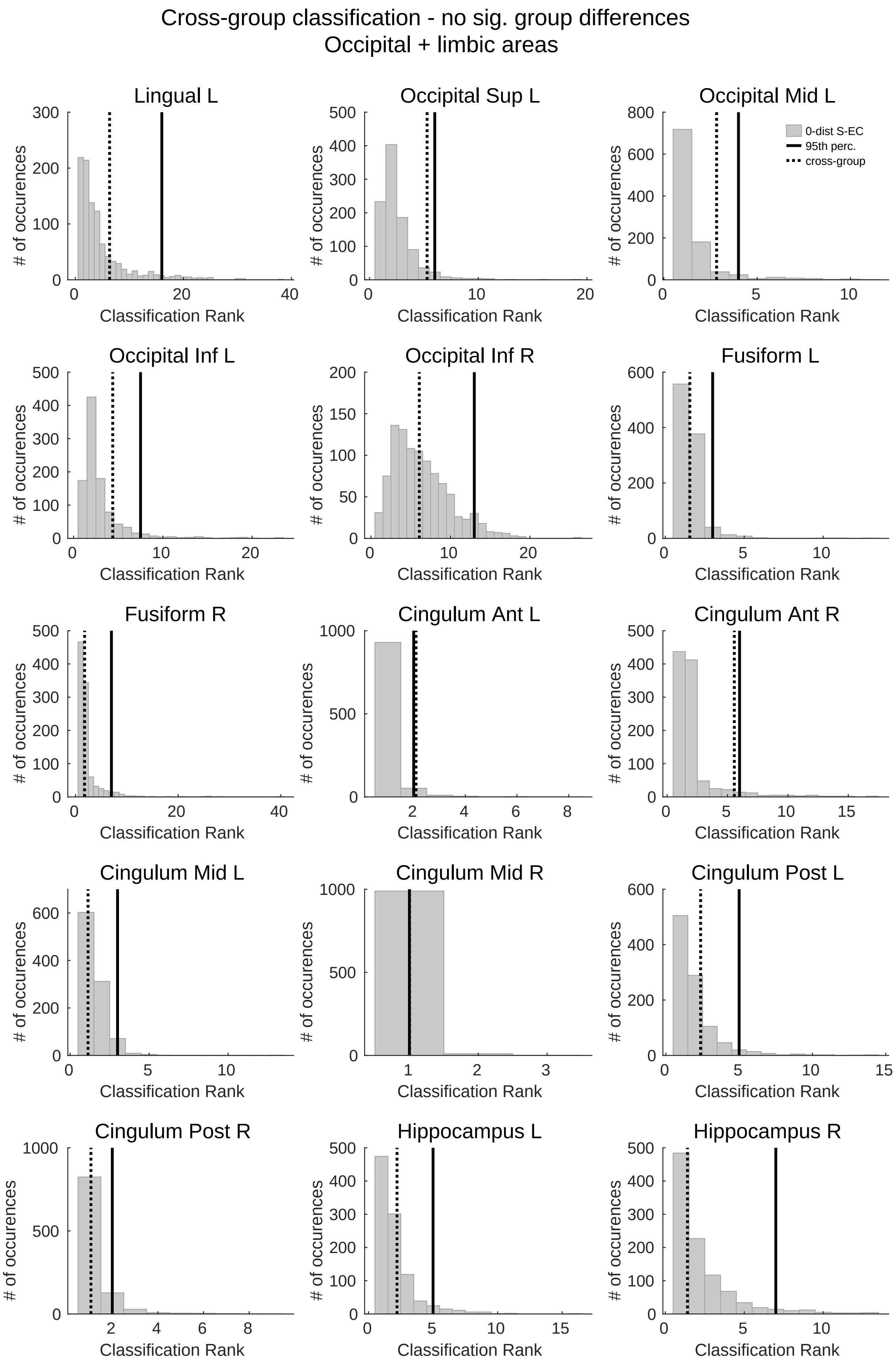

Cross-group classification - no sig. group differences
Occipital + limbic areas
Lingual L
Occipital Sup L
Occipital Mid L
0-dist S-EC
95th perc.
cross-group
Occipital Inf L
Occipital Inf R
Fusiform L
Fusiform R
Cingulum Ant L
Cingulum Ant R
Cingulum Mid L
Cingulum Mid R
Cingulum Post L
Cingulum Post R
Hippocampus L
Hippocampus R
### of occurences
Classification Rank

Cross-group classification - no sig. group differences
Limbic areas

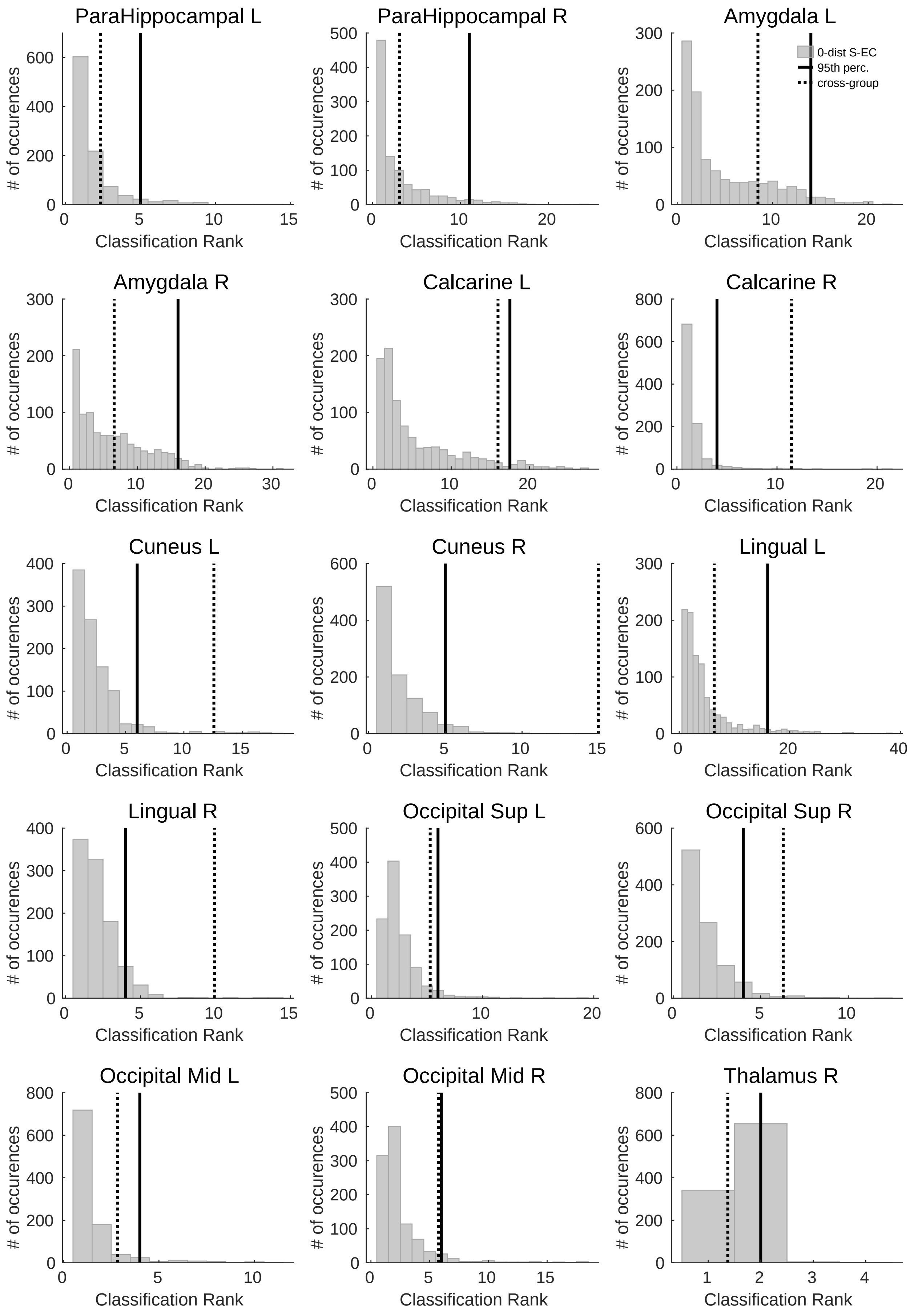

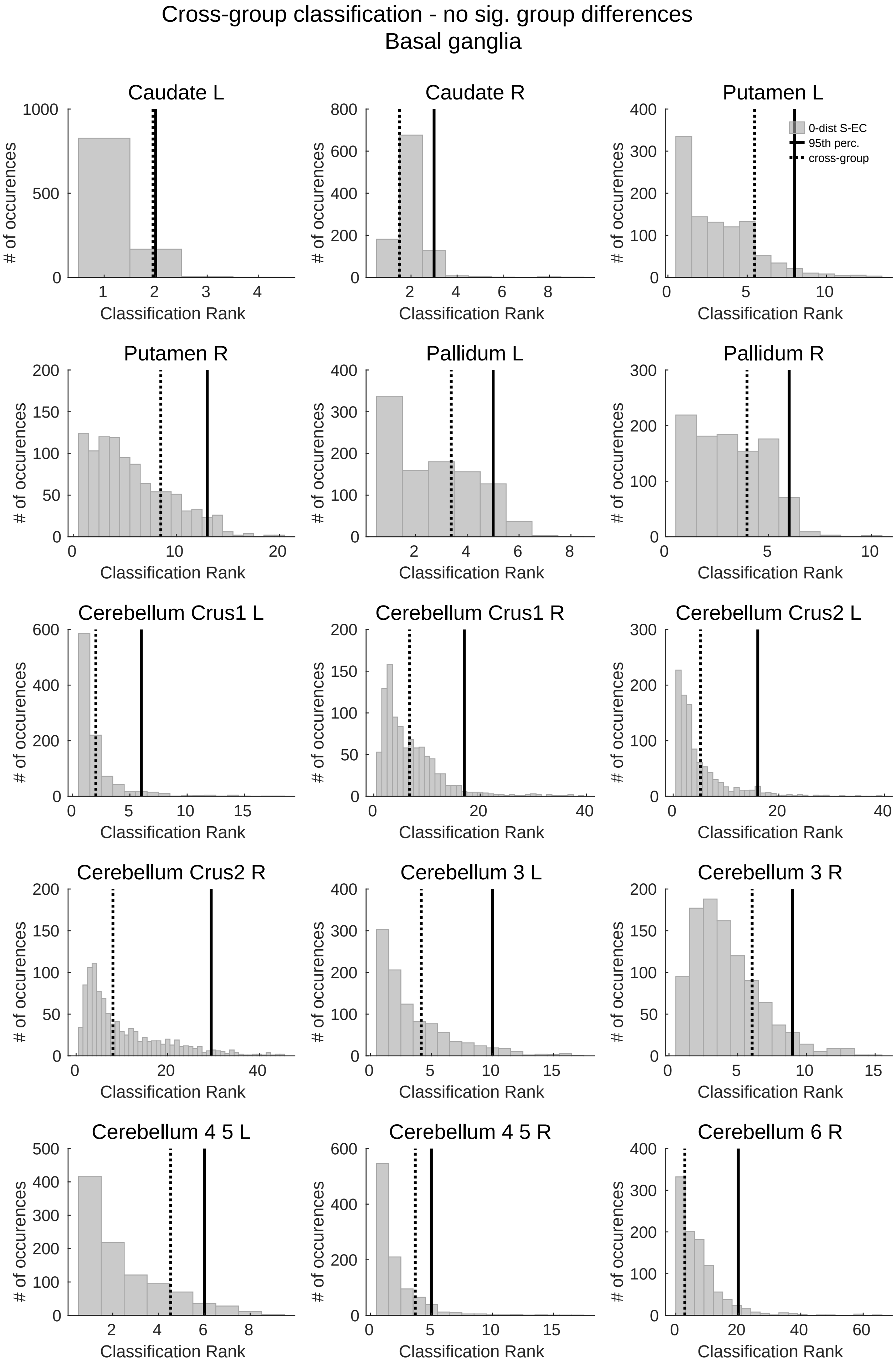

Cross-group classification - no sig. group differences
Basal ganglia
Caudate L
Caudate R
Putamen L
Putamen R
Pallidum L
Pallidum R
Cerebellum Crus1 L
Cerebellum Crus1 R
Cerebellum Crus2 L
Cerebellum Crus2 R
Cerebellum 3 L
Cerebellum 3 R
Cerebellum 4 5 L
Cerebellum 4 5 R
Cerebellum 6 R
0-dist S-EC
95th perc.
cross-group
### of occurences
Classification Rank

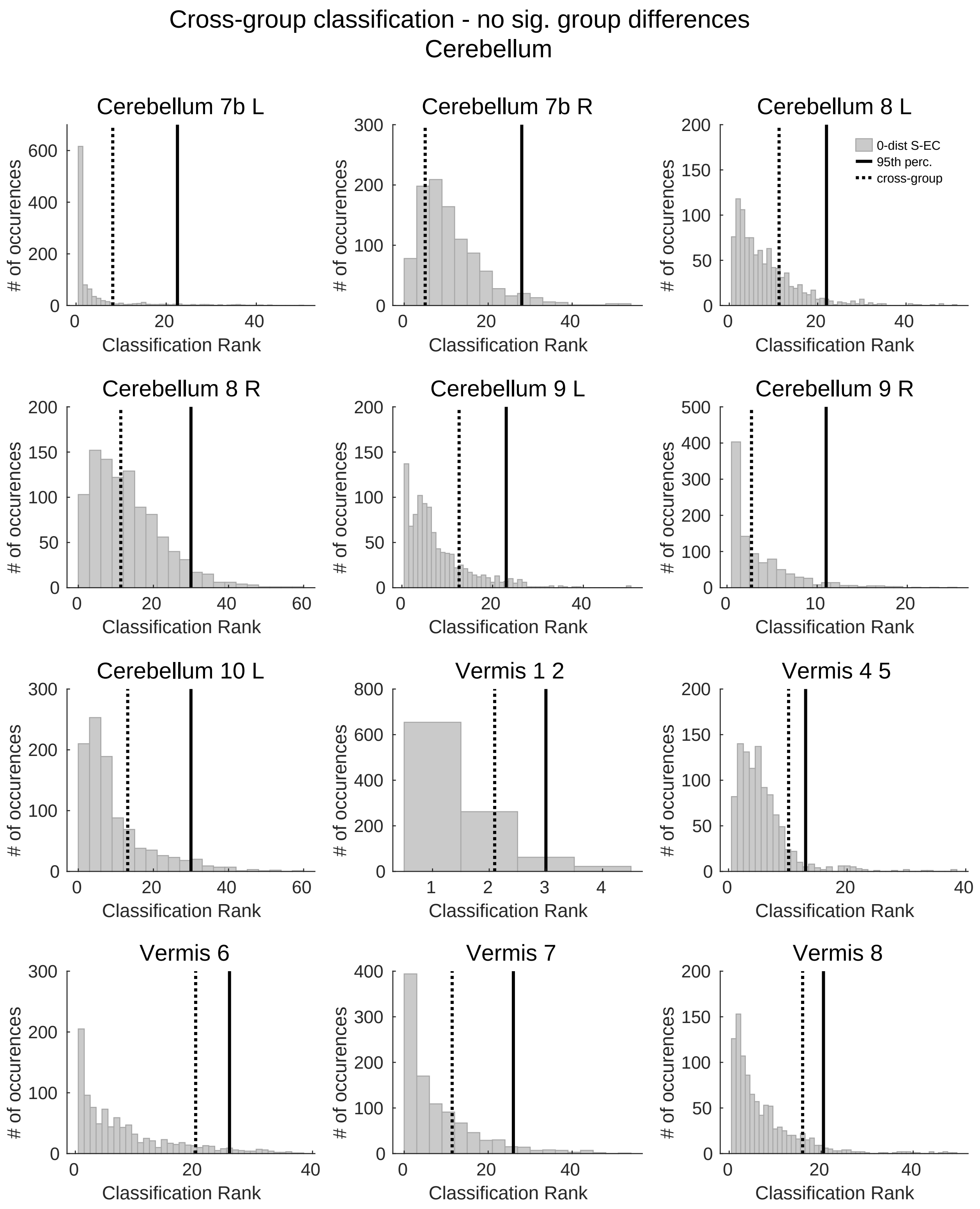

Cross-group classification - no sig. group differences
Cerebellum
Cerebellum 7b L
Cerebellum 7b R
Cerebellum 8 L
0-dist S-EC
95th perc.
cross-group
Cerebellum 8 R
Cerebellum 9 L
Cerebellum 9 R
Cerebellum 10 L
Vermis 1 2
Vermis 4 5
Vermis 6
Vermis 7
Vermis 8
### of occurences
Classification Rank

**Fig. S2. Region-specific distribution of sampled ranks overlaid by cross-group mean rank.** To test whether cross-group classification ranks were distinct from the control condition (S-EC), cross-group ranks were tested against a distribution of ranks from the S-EC. Distributions, generated from classification ranks of corresponding brain areas (i.e. Calcarine in training and test group) across 1000 iterations, are displayed individually for all brain areas (N=115). It was tested whether the cross-group mean rank exceeded the 95$^{th}$ percentile of the null-distribution. Each panel represents one brain area. The 95$^{th}$ percentile is shown as solid line and cross-group mean rank as dotted line.

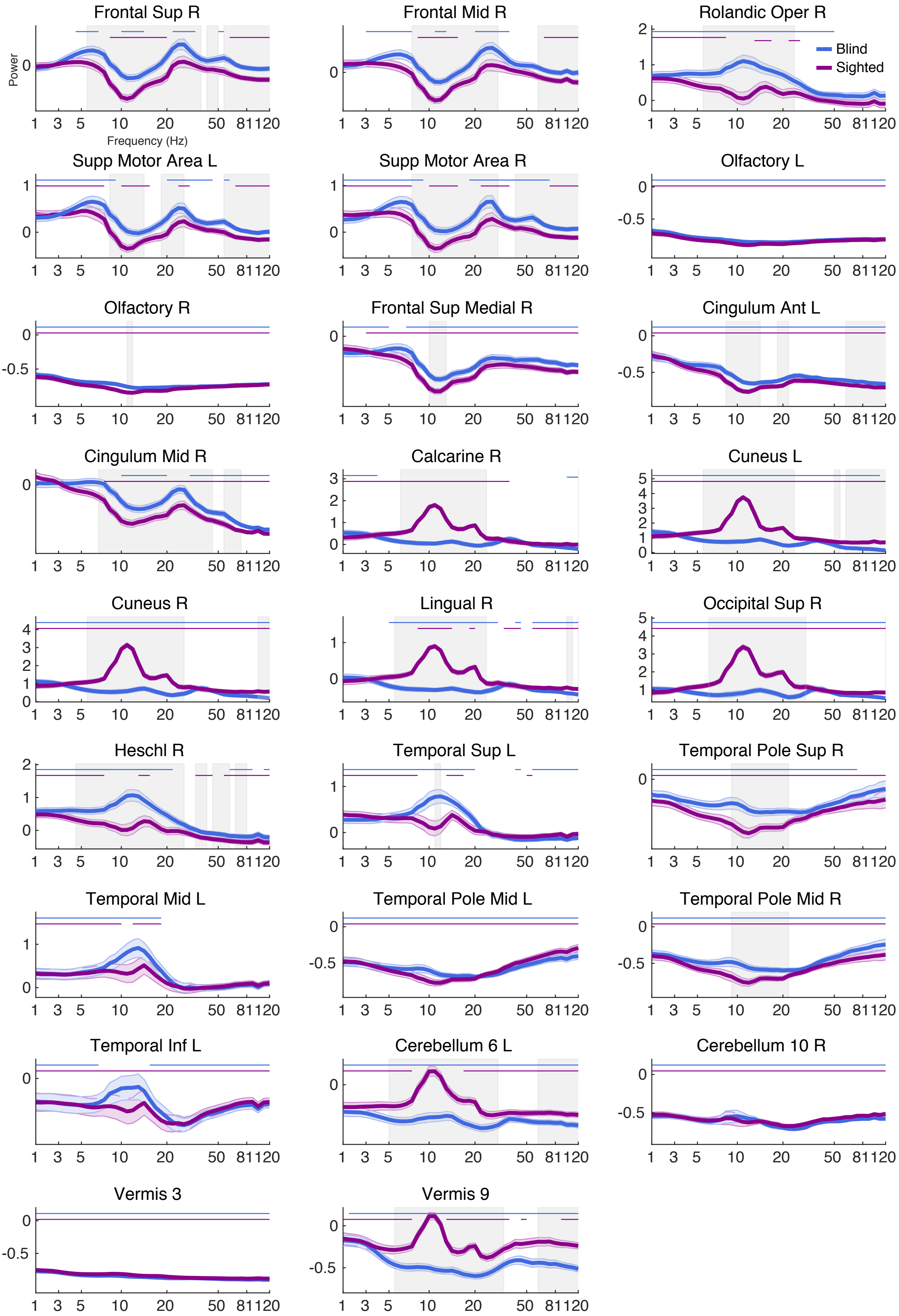
Frontal Sup R
Frontal Mid R
Rolandic Oper R
Blind
Sighted
Power
Frequency (Hz)
1 3 5 10 20 50 81120
Supp Motor Area L
Supp Motor Area R
Olfactory L
Olfactory R
Frontal Sup Medial R
Cingulum Ant L
Cingulum Mid R
Calcarine R
Cuneus L
Cuneus R
Lingual R
Occipital Sup R
Heschl R
Temporal Sup L
Temporal Pole Sup R
Temporal Mid L
Temporal Pole Mid L
Temporal Pole Mid R
Temporal Inf L
Cerebellum 6 L
Cerebellum 10 R
Vermis 3
Vermis 9

**Fig. S3. Post-hoc analysis of spectral differences between S-EC and CB.** Raw normalized spectral profiles (i.e. no clustering) for all brain regions for which cross-group classification was significantly worse compared to classification in the S-EC. Each panel illustrates spectral power for the S-EC (magenta) and the CB (blue), averaged across subjects of each group. Shaded error bars reflect standard error of the mean across subjects. Frequency areas shaded in grey indicate frequency ranges in which the groups showed significant power differences, as assessed by permutation statistics (Q = 0.05; FDR corrected $p$-value = .033; $p$-values < .033). Additionally, we statistically tested spectral power in all depicted ROIs against a baseline of zero at each frequency bin between 1-120 Hz. Due to the ratio normalization, testing power peaks against "zero" here indicates whether at a certain ROI the power is greater compared to the average power across the brain at this frequency. Significant frequencies are illustrated by horizontal lines separately for the groups. The figure demonstrates that in many areas spectral power was significantly different from "zero" across a broad range of frequencies, and crucially for all frequency bands we discuss in the manuscript.

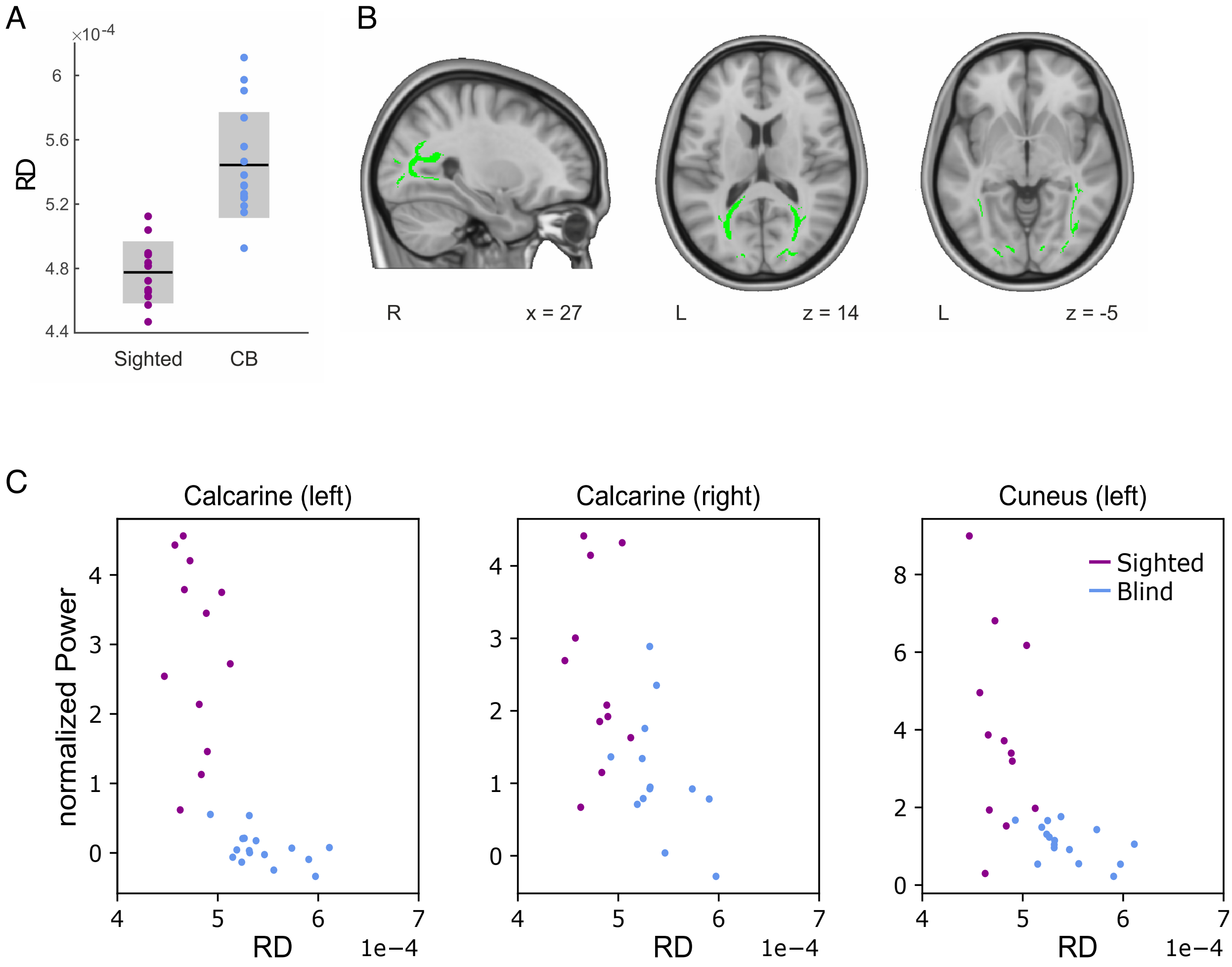

A
B
C
×10-4
RD
Sighted
CB
R
x = 27
L
z = 14
L
z = -5
Calcarine (left)
Calcarine (right)
Cuneus (left)
normalized Power
RD
1e−4
Sighted
Blind

**Fig. S4. Analysis of microstructural properties. A)** Anatomical connectivity data indicated structural group differences between the congenitally blind and sighted groups. Radial diffusivity (RD) statistical contrast maps were generated using tract-based spatial statistics (TBSS) and display congenitally blind over sighted participants' values. To facilitate the visualization of the pattern of results, box plots ($N_{CB}$ = 16, $N_{Sighted}$ = 12) with the mean (center line) ± *SD* (gray area) RD values of the significant cluster are shown for each group. **B)** Images of the white matter cluster (green) that showed significant group differences ($N_{CB}$ = 16, $N_{Sighted}$ = 12; family-wise error (FWE) corrected, two-sided $p < .05$, threshold- free cluster enhancement) are shown. Neurological convention is used, with MNI coordinates at the bottom of each slice. **C)** Scatterplots visualize relations between region (left and right calcarine, left cuneus) and frequency specific (alpha) spectral clusters and RD values. More specifically, spectral clusters for which correlations (Spearman correlation, FDR-corrected: Q = 0.05, $p < .0065$) between RD and power were significant, are illustrated (cf. Table S1 for the results of all correlations and a more detailed description of the correlation analysis). Dots represent individual subjects and are color-coded according to the group (blue = congenitally blind, magenta = sighted).

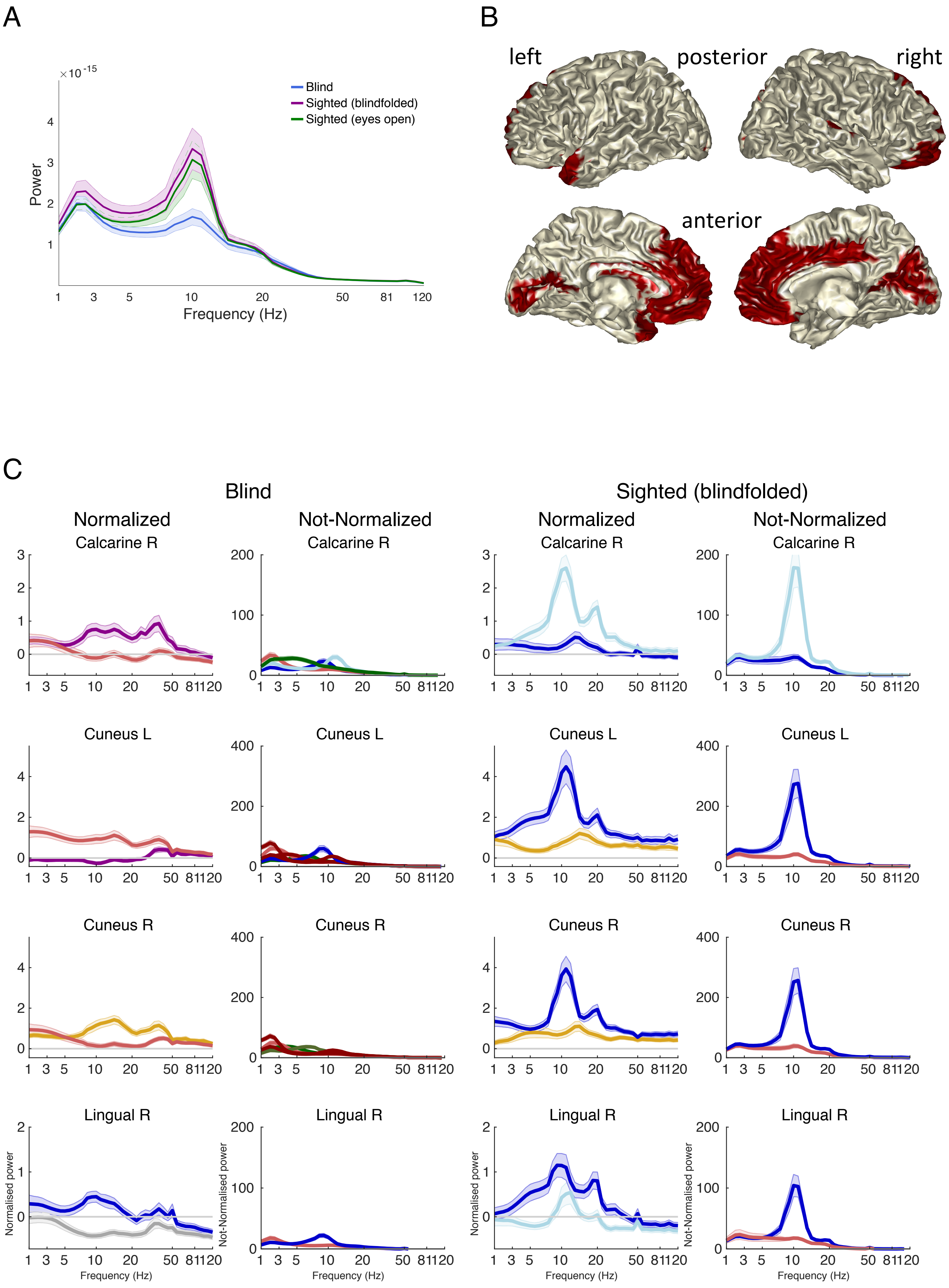

A
B
C
×10^-15
Power
Frequency (Hz)
Blind
Sighted (blindfolded)
Sighted (eyes open)
left
posterior
right
anterior
Blind
Sighted (blindfolded)
Normalized
Not-Normalized
Calcarine R
Cuneus L
Cuneus R
Lingual R
Normalised power
Not-Normalised power

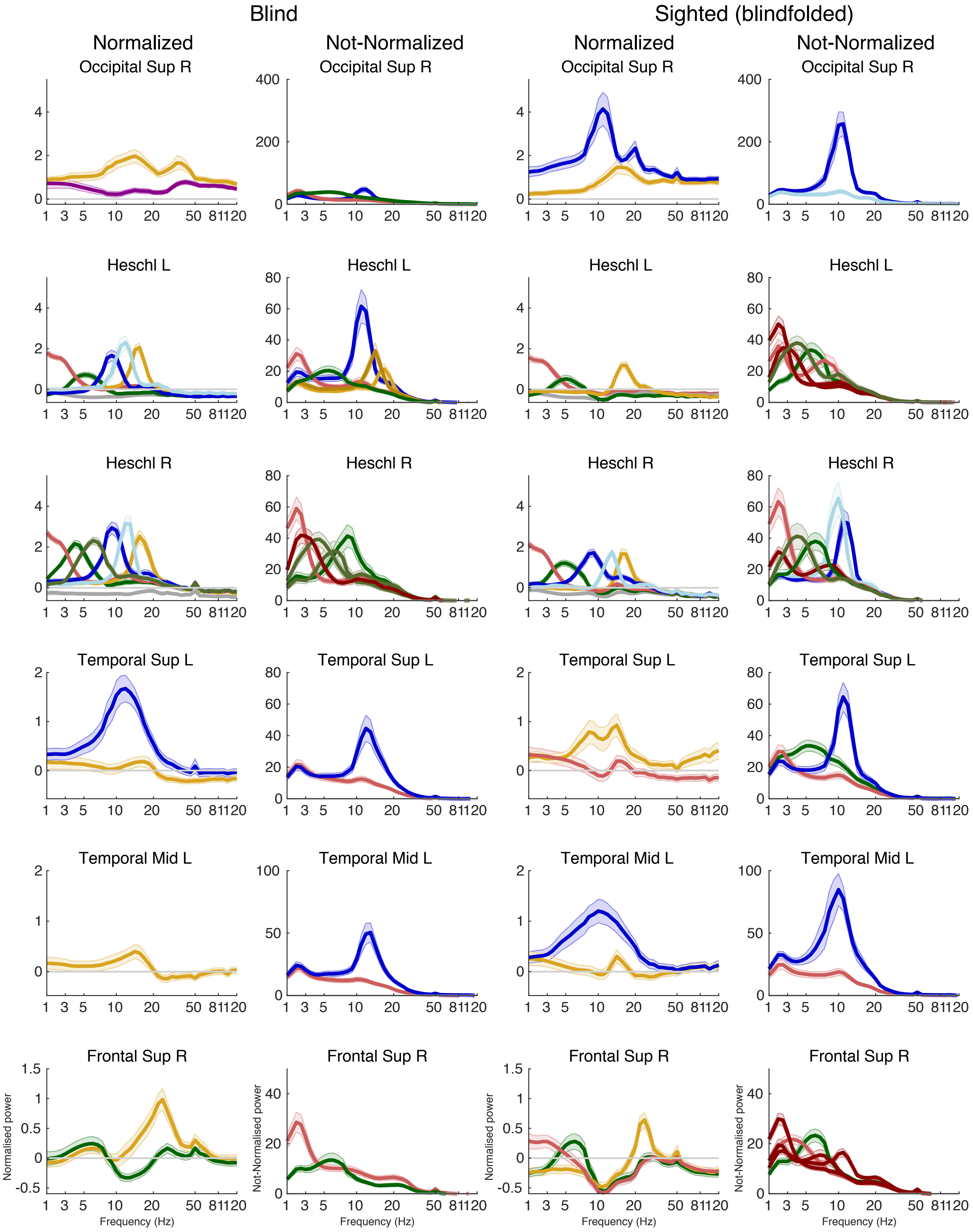
Blind
Sighted (blindfolded)
Normalized
Not-Normalized
Occipital Sup R
Heschl L
Heschl R
Temporal Sup L
Temporal Mid L
Frontal Sup R
Normalised power
Not-Normalised power
Frequency (Hz)

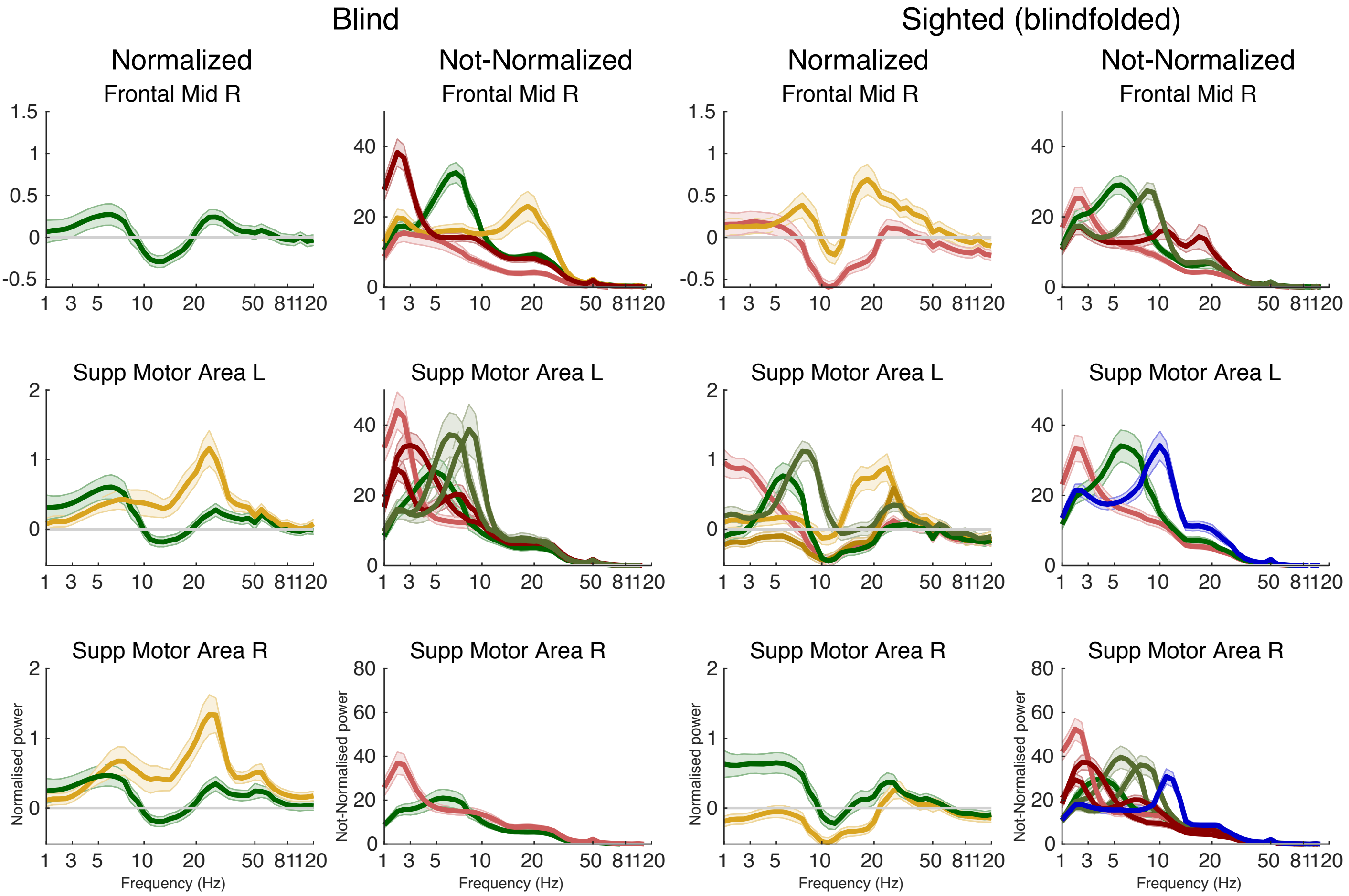

Blind
Sighted (blindfolded)
Normalized
Not-Normalized
Normalized
Not-Normalized
Frontal Mid R
Supp Motor Area L
Supp Motor Area R
Normalised power
Not-Normalised power
Frequency (Hz)

**Fig. S5. Normalization spectrum. A)** Prior to the clustering procedure, Fourier spectra were ratio normalized (i.e. per participant, power at each voxel and segment was divided by the average power across temporal segments and voxels). The ratio normalization has the advantage of removing the predominant neuronal activity (in the alpha and super-low frequency ranges) from the power spectrum known to otherwise hinder the spectral analysis of lower frequency neuronal activity. Here, the average power spectrum across segments and voxels is depicted, averaged across participants. Shaded error bars reflect the standard error of the mean across participants. The x-axis illustrates frequency, scaled logarithmically from 1 to 120 Hz; the y-axis illustrates power. For visualization, power was interpolated at 50 Hz due to line noise. Each line corresponds to one of the experimental groups (blue: congenitally blind, magenta: sighted (eyes closed), green: sighted (blindfolded)). **B)** In order to test the impact of the normalization procedure on our group findings, here we performed the clustering procedure and the cross-group classification on the non-normalized data. The cross-group classification analysis confirmed our main findings for the normalized data, by showing significantly worse classification in visual, temporal and right frontal areas compared to a null-distribution (classification within the sighted; bilateral: superior frontal gyrus (medial), medial orbital frontal gyrus, gyrus rectus, anterior cingulate gyrus, calcarine gyrus; right: superior frontal gyrus (orbital), middle frontal gyrus (orbital), middle cingulate gyrus, cuneus, Heschl gyrus; left: caudate nucleus, superior temporal pole, middle temporal pole, olfactory cortex). However, with the non-normalized data overall less brain areas showed significant effects. **C)** Clustered spectral profiles derived from the non-normalized data shown separately for a selection of brain areas (rows) for the congenitally blind (first and second column) and blindfolded sighted participants (third and fourth columns). The first and the third columns represent normalized data, while the second and fourth columns show non-normalized data. The comparison of the spectral profiles of the normalized and the non-normalized spectra displays two observations: First, overall there was congruence between the normalized and non-normalized spectra; Second, the normalized spectra showed a reduced presence of the strong alpha and low-frequency power across the brain, demonstrating that our normalization procedure was successful.

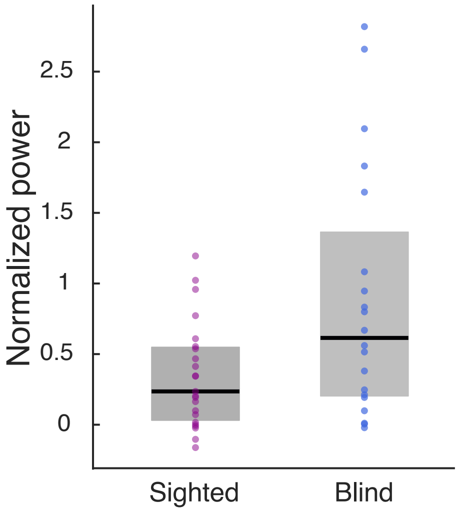


**Fig. S6. Statistical comparison of gamma band power in Calcarine gyrus in the congenitally blind and sighted.** In the congenitally blind individuals, the clustering procedure revealed a spectral cluster with a power peak in the low gamma frequency band in right Calcarine gyrus. In contrast, no power peak was apparent in the spectral profile of the sighted individuals in the same frequency band and brain area. When the unclustered spectra were statistically compared (cf., post-hoc analysis in S4), we found no significant group differences within the low gamma band. The comparison of the unclustered spectra is a more conventional approach, which however, is lacking the advantages of the clustering procedure. The clustering procedure allows to detect more nuanced spectral effects by separating distinct spectral clusters. Thus, to further investigate the apparent group difference in gamma power in right Calcarine gyrus, we statistically compared the gamma power as obtained from the clustering procedure. To this end, all single-subject clusters contributing to the group cluster in right Calcarine gyrus were identified. For subjects contributing multiple $1^{st}$-level clusters to the group cluster, $1^{st}$-level clusters were averaged which resulted in one cluster per subject. Peak power within the low gamma band (33.5-50 Hz) was extracted and a Mann-Whitney-U-test was performed to test for statistical group differences. The analysis showed higher gamma power in the congenitally blind individuals compared to the sighted ($z = -2.082$, $p = .0374$).

| | | Alpha band | | Gamma band | |
|---|---|---|---|---|---|
| **ROI** | **H** | **rho** | ***p*** | **rho** | ***p*** |
| Calcarine | L | -0.81 | < .001* | -0.18 | .371 |
| Calcarine | R | -0.55 | .005* | 0.36 | .083 |
| Cuneus | L | -0.67 | < .001* | -0.09 | .666 |
| Cuneus | R | -0.29 | .13 | 0.24 | .222 |
| Lingual | L | 0.57 | .007 | 0.40 | .070 |
| Lingual | R | 0.01 | .960 | -0.08 | .727 |
| SOG | L | -0.51 | .022 | 0.15 | .535 |
| SOG | R | -0.19 | .333 | 0.46 | .016 |
| MOG | L | 0.38 | .049 | 0.43 | .026 |
| MOG | R | 0.02 | .913 | 0.37 | .057 |
| IOG | L | -0.52 | .013 | 0.15 | .493 |
| IOG | R | -0.5 | .015 | 0.16 | .480 |

**Table S1. Correlation between RD values and spectral power.** For the correlation analysis, all brain areas in occipital cortex were included (if power peaks were clearly distinguishable). For each of these brain regions the power peaks of clusters were retrieved in the following way: the single subject clusters (i.e., first-level clusters) that contributed to each cluster of a spectral profile on the group-cluster level were identified and individual power peaks were extracted. Power values were correlated with RD values (Spearman correlation, FDR-corrected: Q = 0.05, corrected $p$-value = .0065, $p$-values < .0065). ROI - region of interest; H - hemisphere; IOG – inferior occipital gyrus; L - left; MOG - middle occipital gyrus; R - right; SOG - superior occipital gyrus.
